# Convergent evolution of (βα)_8_-barrel fold methylene-tetrahydropterin reductases utilizing a common catalytic mechanism

**DOI:** 10.1101/2023.09.18.558202

**Authors:** Manuel Gehl, Ulrike Demmer, Ulrich Ermler, Seigo Shima

**Affiliations:** Max Planck Institute for Terrestrial Microbiology, Marburg, Germany; Max Planck Institute of Biophysics, Frankfurt am Main, Germany

**Keywords:** crystal structure, evolution, methylene-tetrahydrofolate reductase, methylene-tetrahydromethanopterin reductase, catalytic mechanism

## Abstract

Methylene-tetrahydropterin reductases are folded in (βα)_8_ barrel and catalyze the reduction of a methylene to a methyl group bound to a reduced pterin as C_1_ carrier in various one-carbon (C_1_) metabolisms. F_420_-dependent methylene-tetrahydromethanopterin (methylene-H_4_MPT) reductase (Mer) and the flavin-independent methylene-tetrahydrofolate (methylene-H_4_F) reductase (Mfr) use a ternary complex mechanism for the direct transfer of a hydride from F_420_H_2_ and NAD(P)H to the respective methylene group, whereas FAD-dependent methylene-H_4_F reductase (MTHFR) uses FAD as prosthetic group and a ping-pong mechanism to catalyze the reduction of methylene-H_4_F. A ternary complex structure of MTHFR is available and based on this structure, a catalytic mechanism was proposed, while no ternary complex structures of Mfr or Mer are reported. Here, Mer from *Methanocaldococcus jannaschii* (jMer) was heterologously produced and the crystal structures of the enzyme with and without F_420_ were determined. A ternary complex of jMer was modeled using a functional alignment approach based on the ternary complex structure of MTHFR and the modeled ternary complex of Mfr. Mutational analysis at the structurally conserved positions of the three reductases indicated that although these reductases share a limited sequence identity, the key catalytic glutamate residue is conserved and a common catalytic mechanism involving the formation of a 5-iminium cation of the methylene-tetrahydropterin intermediate is shared. A phylogenetic analysis indicated that the three reductases do not share one common ancestor and the conserved active site structures of the three reductases may be the result of convergent evolution.

**STATEMENT:** This work provides evidence for a common catalytic mechanism of the functional class of methylene-tetrahydropterin reductases. Despite their very low sequence identity, they share a (βα)_8_-barrel structure with a similar active site geometry. Phylogenetic and mutational analyses suggested that these enzymes have developed from distinct ancestors as a result of convergent evolution. This work describes an example of a catalytic mechanism that emerged independently for several times during evolution in the three domains of life.

## INTRODUCTION

Redox reactions of C_1_ units bound to C_1_ carriers are widespread in the three domains of life. The most common C_1_ carriers are tetrahydrofolate (H_4_F) and tetrahydromethanopterin (H_4_MPT), which consist of a reduced pterin (tetrahydropterin) bound to a *para*-aminobenzoate (PABA) group and a variable tail region [1]. The bound C_1_ unit is transformed between oxidation states of formic acid (+II, formyl and methenyl groups), formaldehyde (0, methylene group) and methanol (−II, methyl group) in the C_1_ metabolisms. Examples for such metabolisms are the methanogenic pathways, the methyl-branch of the Wood-Ljungdahl pathway, and the folate cycle [2,3].

The reduction of methylene-H_4_F to methyl-H_4_F is catalyzed by methylene-H_4_F reductases [3–5], which are classified into a flavin-dependent enzyme (MTHFR) and a flavin-independent one (Mfr) [5,6]. MTHFR is further classified into subclasses depending on either FAD or FMN as a prosthetic group [4,7–10].

The FAD-dependent MTHFR uses a ping-pong reaction mechanism (Figure 1A), in which the tightly bound FAD is first reduced by NAD(P)H and then FADH_2_ reduces methylene-H_4_F [11]. The structural characterization of the inactive MTHFR_Glu28Gln mutant from *E. coli* (eMTHFR) with FAD and with either NADH or methyl-H_4_F [12] revealed both substrates in the same position, explaining the structural basis for the ping-pong mechanism. Since the imidazolidine ring in methylene-H_4_F is a poor acceptor for the negatively charged hydride, the substrate is activated to a 5-iminium cation [13]. A 5-iminium cation intermediate was also proposed in the non-enzymatic condensation of formaldehyde and H_4_F to form methylene-H_4_F [14]. Additionally, in the crystal structure of thymidylate synthase, a 5-hydroxymethylene-H_4_F was found, which supports the presence of the 5-iminium cation as an intermediate in this enzyme reaction [15]. Based on the drastic effect of the Glu28Gln mutation of eMTHFR on the activity, Glu28 was proposed to be a catalytic base for the formation of the 5-iminium cation.

**Figure 1:**
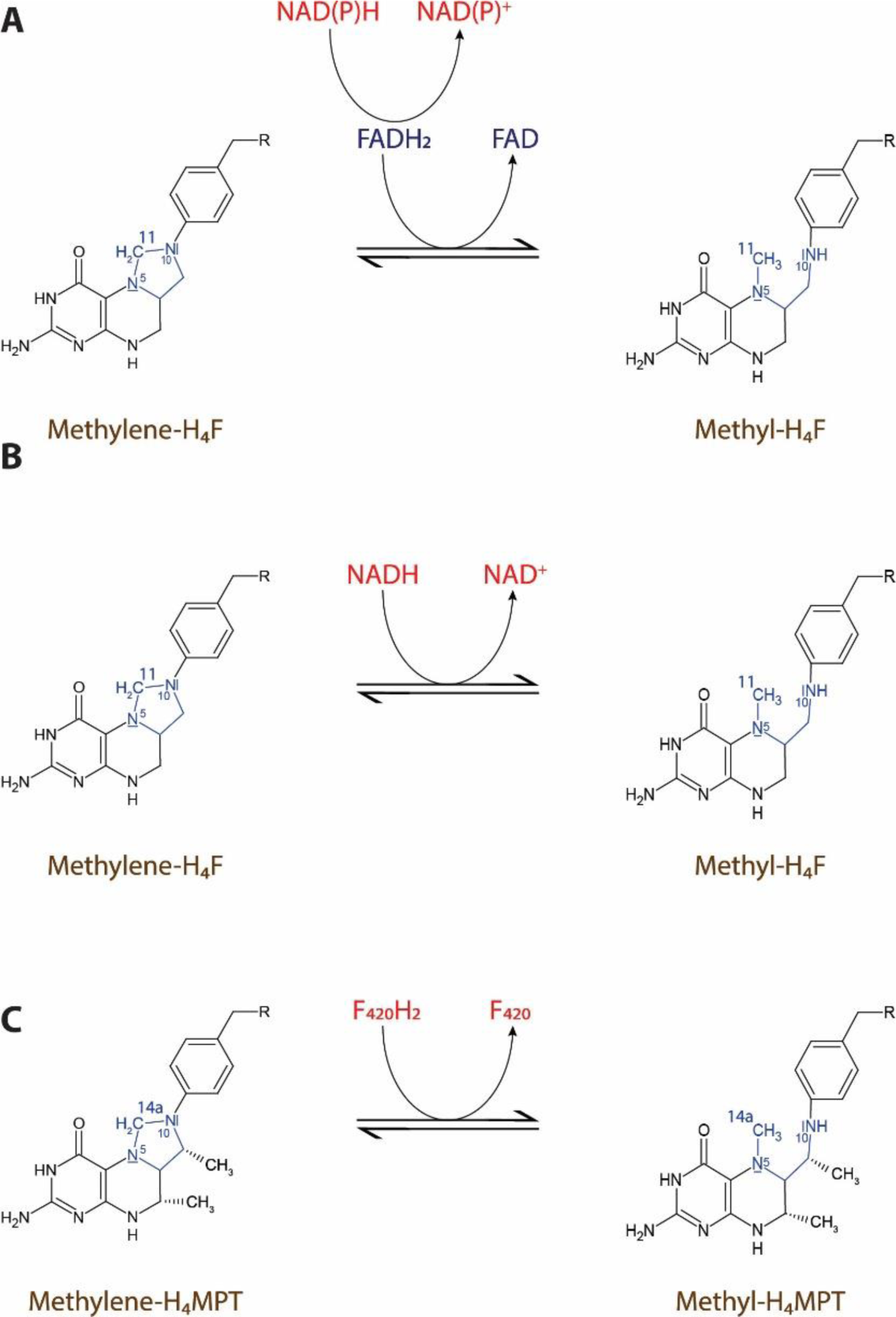
Reactions catalyzed by methylene-tetrahydropterin reductases. (A) FAD-dependent methylene-H_4_F reductases (MTHFR) catalyzes the reduction of methylene-H_4_F using reduced FADH_2_ as reductant. FAD is bound to the enzyme as a prosthetic group and is reduced first by NAD(P)H. (B) Flavin-independent methylene-H_4_F reductase (Mfr) catalyzes the reduction of methylene-H_4_F using NADH as reducing agent. In contrast to MTHFR, Mfr has no prosthetic group and uses a ternary complex reaction mechanism rather than a ping-pong mechanism. (C) F_420_-dependent methylene-H_4_MPT reductase (Mer) catalyzes the reduction of methylene-H_4_MPT using reduced F_420_ (F_420_H_2_) as reducing agent.

Mfr is a monomeric enzyme found only in mycobacteria. Recently, the enzymes encoded by *MSMEG_6596* and *MSMEG_6649* in *Mycobacterium smegmatis* were found to have MTHFR activity [5]. Both enzymes do not contain flavins and catalyze a direct hydride transfer from NADH to methylene-H_4_F by a ternary complex mechanism (Figure 1B). A knockout strain of *M. smegmatis* showed impaired growth in the absence of methionine, suggesting that Mfr is involved in the methionine cycle in mycobacteria [5]. Likewise, an Mfr homolog encoded by *Rv2172c* in *M. tuberculosis* is essential for the growth of this organism [6]. Recently, we reported the crystal structure of Mfr from *Mycolicibacterium hassiacum* (hMfr) and gave evidence that Glu9 in the active site is the key catalytic residue and that the 5- iminium cation forming mechanism of eMTHFR and hMfr are identical.

Most methanogenesis pathways involve the methylene-H_4_MPT reductase (Mer), which catalyzes the reversible reduction of methylene-H_4_MPT with F_420_H_2_ as reductant using a ternary complex mechanism [16,17] (Figure 1C). Crystal structures are available for substrate-free Mer from *Methanopyrus kandleri* and *Methanothermobacter marburgensis* [18] and the *Methanosarcina barkeri* Mer-F_420_ binary complex [19]. Mer belongs to the bacterial luciferase family, which consists of FMN- and F_420_-dependent oxidoreductases [19].

Here, we report the crystal structure of Mer from *Methanocaldococcus jannaschii* (jMer) with and without F_420_. Using a functional structural alignment, we identified similar amino acid residues at equivalent positions in the active sites of eMTHFR, jMer and hMfr, which are possibly involved in the binding of the C_1_ carrier, and in activation of the substrate by protonation. We investigated the role of specific amino acids by mutational and kinetic analysis. Evidence has been provided that all three methylene-tetrahydropterin reductases have the same fold, a similar methylene-tetrahydropterin binding position, and the same catalytic mechanism, although the binding mode of the C_1_ carriers and reductants is not identical.

## RESULTS AND DISCUSSION

### Characterization of the heterologously produced jMer

Attempts to produce Mer from different organisms in *E. coli* have failed in the past because Mer formed inclusion bodies [20]. However, recently, the first successful heterologous expression of Mer from *Methanocaldococcus jannaschii* (jMer) was reported, in which the heterologously produced jMer catalyzed the formation of lactaldehyde by reduction of methylglyoxal with NADPH [21]. We obtained the published expression system from the authors and characterized the purified enzyme. The *K*_m_ and *V*_max_ values of jMer are in the range of those of the respective enzymes from methanogenic archaea (*M. barkeri*, *M. marburgensis*, and *M. kandleri*) and sulfate-reducing archaea (*Archaeoglobus fulgidus*) (Table 1) [16,22–25]. The comparison of the catalytic constants of jMer, eMTHFR and hMfr shows that the reaction rate of jMer is about 30 times higher than that of hMfr and eMTHFR (Table 2). This finding can be explained as eMTHFR and hMfr are involved in anabolism while jMer in catabolism, which requires higher enzymatic turnover rates. Size-exclusion chromatography showed that jMer in solution is a homodimer (∼80 kDa) (Figure S1) as reported in the literature [21].

**Table 1:** Comparison of the catalytic constants of Mer from *Methanosarcina barkeri*, *Methanothermobacter marburgensis*, *Methanopyrus kandleri*, *Archaeoglobus fulgidus*, *Methanocaldococcus jannaschii*.

| Mer from | | Apparent $K_m$ | Apparent $V_{max}$ | Source |
| --- | --- | --- | --- | --- |
| *M. barkeri* | methylene-$H_4$MPT | 15 μM | 2200 U/mg | [22] |
| | $F_{420}H_2$ | 12 μM | | |
| *M. marburgensis* | methylene-$H_4$MPT | 300 μM | 6000 U/mg | [16] |
| | $F_{420}H_2$ | 3 μM | | |
| *M. kandleri* | methylene-$H_4$MPT | 7 μM | 435 U/mg | [24] |
| | $F_{420}H_2$ | 4 μM | | |
| *A. fulgidus* | methylene-$H_4$MPT | 16 μM | 450 U/mg | [23] |
| | $F_{420}H_2$ | 4 μM | | |
| *M. jannaschii* | methylene-$H_4$MPT | 58 μM | 500 U/mg | This work |
| | $F_{420}H_2$ | 5 μM | | |

**Table 2:**
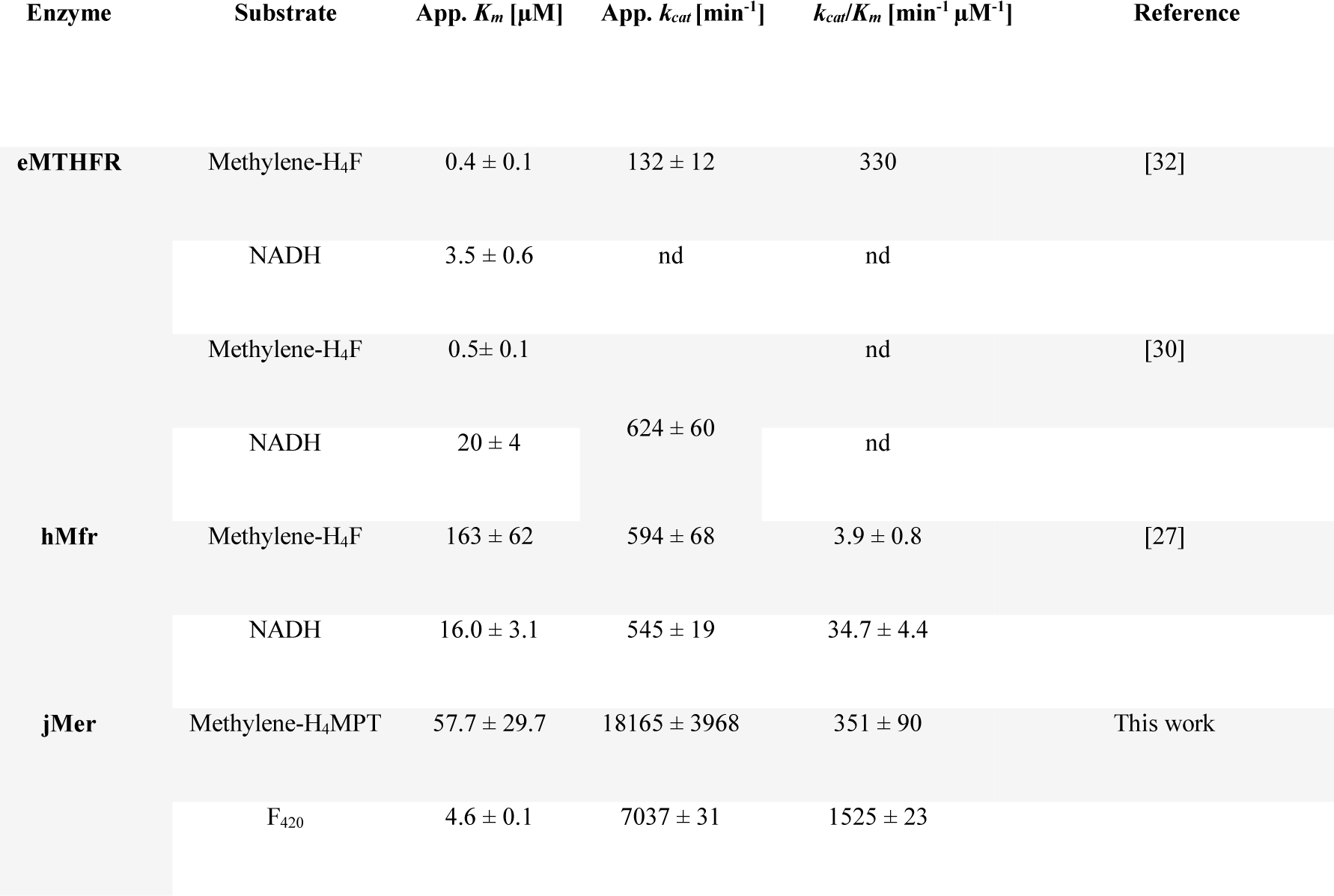
Kinetic constants of eMTHFR, hMfr and jMer. The constants for eMTHFR were obtained from the literature. Where the given *k_cat_* values were not specified with respect to the substrate, the resulting values were placed in the middle of the corresponding row. nd = not determined.

### Determination of the crystal structure of jMer

Crystal structures for the apoenzyme of jMer and the binary jMer-F_420_ complex were determined at a resolution of 1.8 Å and 1.9 Å, respectively (Table 2). The overall structure of jMer is almost identical to those of previously established methanogenic enzymes characterized by a (βα)_8_- or TIM-barrel fold (Figure 2). The same fold was also found in the structures of MTHFR and Mfr [26,27]. The root mean square deviation (RMSD) values of these enzymes are 4.7 Å between eMTHFR and hMfr (over 240 amino acids), 4.6 Å between hMfr and jMer (over 232 amino acids), and 5.3 Å between jMer and eMTHFR (over 200 amino acids).

**Figure 2:**
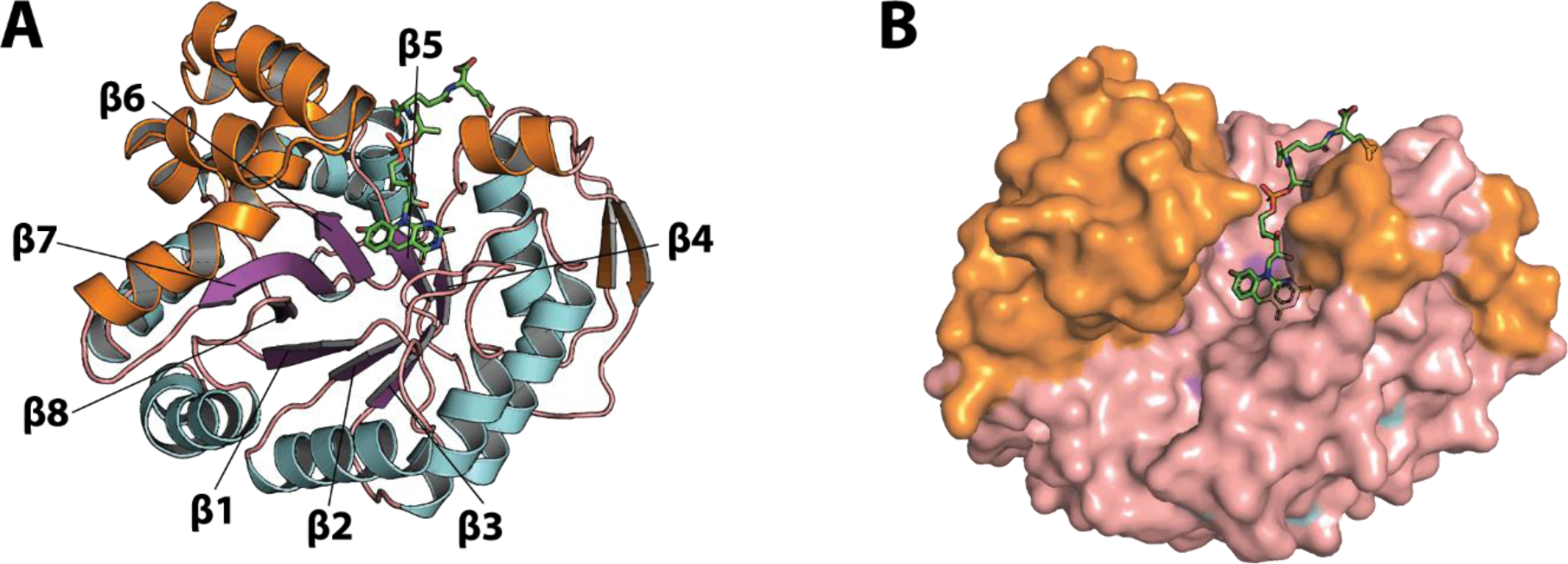
Structure of jMer in complex with F_420_. (A) The β-strands of the (βα)_8_ core unit are labelled and colored purple, while the α-helices of the core unit are colored light blue and the loops are colored salmon. The inserted segments, which form secondary structural elements in addition to the core unit, are colored in orange. F_420_ is shown with carbon in green. (B) The surface model with the stick model of F_420_. The color code is the same as in panel A.

The secondary structural elements are connected by loops typically involved in the active site architecture of TIM barrel enzymes [28,29]. The active site of jMer is located in a cleft between the α/β-barrel domain and a helix-bundle subdomain composed of five helices. In the experimentally determined jMer-F_420_ structure, the cleft is open. F_420_ is associated with the α/β-barrel domain with the isoalloxazine ring positioned at the bottom of the cleft in contact with the C-terminal loops of most strands of the (αβ)_8_ barrel (Figure 2A). The residual part pointing to the entrance of the cleft is sandwiched between the loops after β4 and β5 and reached the N-terminal ends of the following helices (Figure 2B).

The cleft is sufficiently large to also accommodate methylene-H_4_MPT with the pterin head adjacent with the isoalloxazine ring of F_420_. To study the hydride transfer of Mer, we tried to co-crystallize jMer with F_420_/F_420_H_2_ and methylene-/methyl-H_4_MPT. However, the C_1_ carrier could not be found in the resulting electron density. Therefore, we built a model of the ternary complex of jMer for studying the catalytic mechanism.

### Ternary complex model building

To compare the active site structure of eMTHFR, hMfr and jMer, the monomeric protein structures were superposed using a knowledge-based three-dimensional alignment (Figure 3). A simple three-dimensional alignment was not successful because of the large differences between the tertiary structures of jMer and the other two reductases. In this alignment, it was assumed that the hydride transfer process between FADH_2_ and methylene-H_4_F in eMTHFR, and F_420_H_2_ and methylene-H_4_MPT in jMer are the same, resulting in a similar active site geometry. As the first step, the hydride-bearing atoms of FAD (N5) in the eMTHFR structure and of F_420_ (C5) in the jMer structure were superposed (Figure 3, step 2) and then the rest of the proteins was aligned without moving the hydride-transferring atoms (Figure 3, step 3). The hMfr and eMTHFR structures could be superimposed by an overall alignment due to their relatively high structural similarities [27] (Figure 3, step 4). In these aligned tertiary structures, the identical or similar amino-acid residues that locate at the same position in space were identified (Figure 3, step 5). With this aligned models, the position of methyl-H_4_MPT in jMer could be inferred by the coordination of methyl-H_4_F on eMTHFR. The positions of NADH and methyl-H_4_F in hMfr were modeled by us before [27].

**Figure 3:**
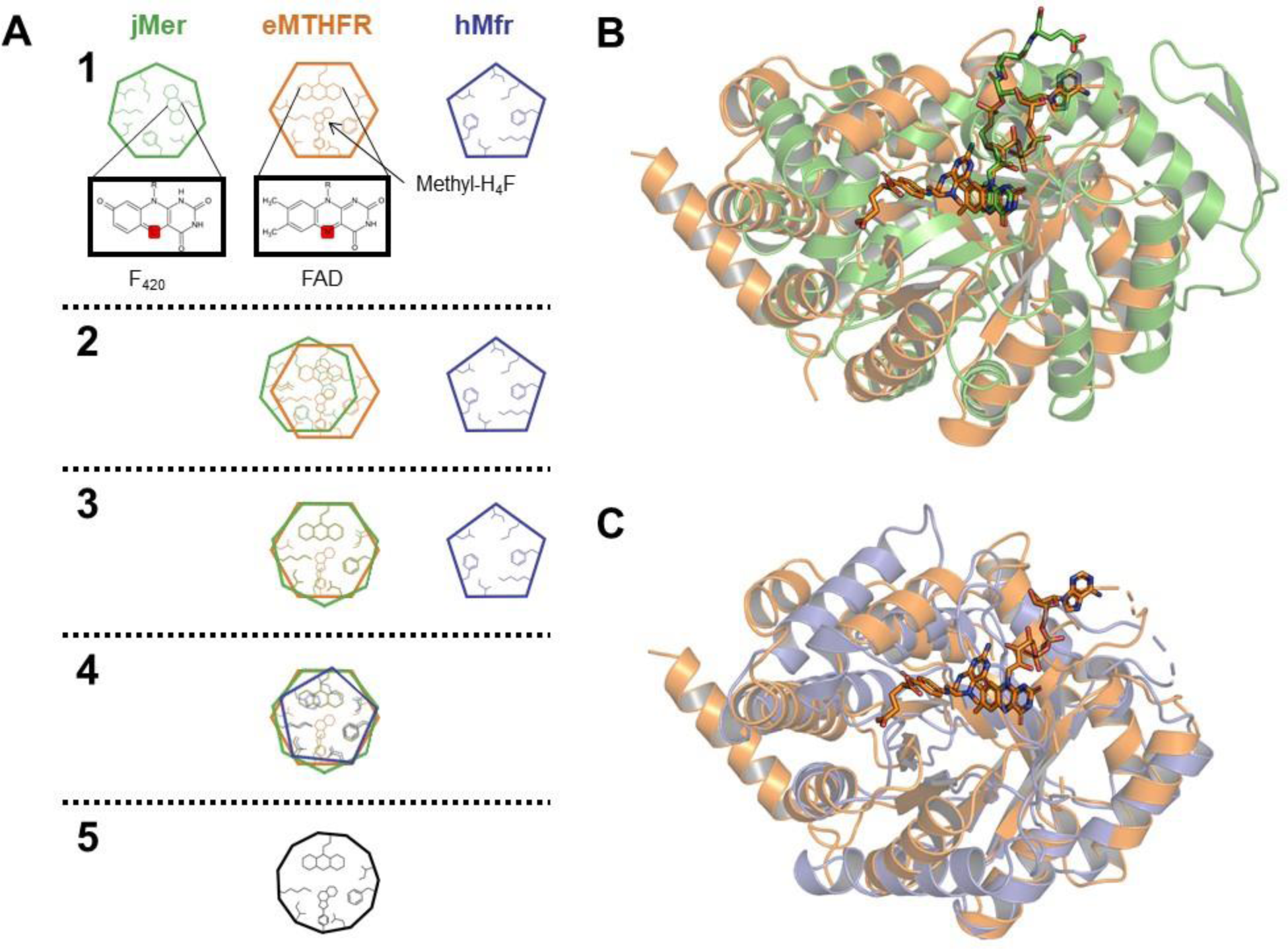
Functional alignment of three methylene-tetrahydropterin reductases. (A) Method of the functional alignment. The hydride-bearing atoms of the reducing agents are highlighted in red (1). N5 of FAD and C5 of F_420_ were aligned (2) and then jMer was aligned on eMTHFR in fixing two hydride-bearing atoms (3). hMfr was aligned to eMTHFR (4). The conserved amino-acid residues were determined based on a structural alignment (5). The result of the alignment of jMer (green) onto eMTHFR (orange) (B), and Mfr (blue) onto eMTHFR (orange) (C). Methyl-H_4_F and FAD of eMTHFR are shown by a stick model with carbon in orange. F_420_ of jMer is shown by a stick model with carbon in green.

In the resulting ternary complex model of jMer, the distance between C11 of the modelled methyl-H_4_F and C5 of F_420_ is 3.3 Å, which is suitable for a hydride transfer (Figure 4). In the open state, methyl-H_4_F does not substantially interfere with amino acid residues of jMer and its tail has no contact with the protein matrix. Notably, there is a hydrophobic pocket consisting of the conserved residues Phe233, Val8 and Val230 near the modelled PABA ring, which might be responsible for binding of the PABA ring of H_4_MPT in jMer. Asn178 is located close to the bottom of the cleft at the C-terminal end of β6, which is presumably hydrogen-bonded with the F_420_ isoalloxazine and the pterin ring. Asp96 protruding from the C-terminal end of β4 is also in contact with the pyrimidine ring of H_4_MPT. Glu6 at the C-terminal end of β1 points towards the deaza-isoalloxazine and pterin rings. The mentioned amino acids Glu6, Phe233, Gln178 and Asp96 involved in binding of the pterin part of the C_1_ carrier have spatial counterparts in MTHFR and hMfr. The four sites are referred to as positions A-D (in Figure 5). Positions A and D are occupied by acidic residues, position C by amino acids with carboxamide groups, and position B by large hydrophobic residues.

**Figure 4:**
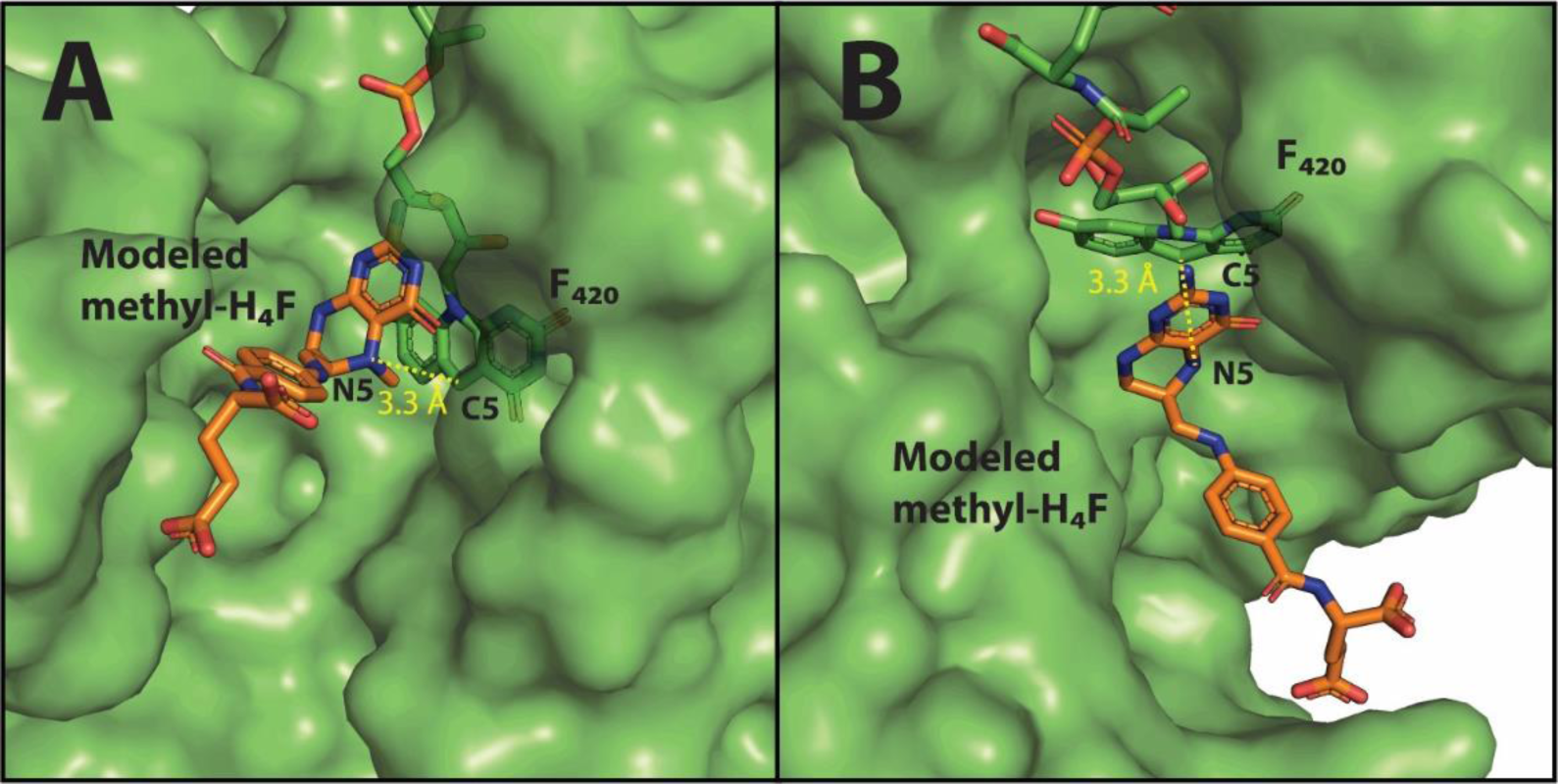
Ternary complex model of jMer built by the functional alignment of the binary complex of jMer and F_420_ (green) and methyl-H_4_F (orange) (see Figure 3). The N5-methyl group of methyl-H_4_F is shown in red. The active-site structure is shown from different angles (A and B).

**Figure 5:**
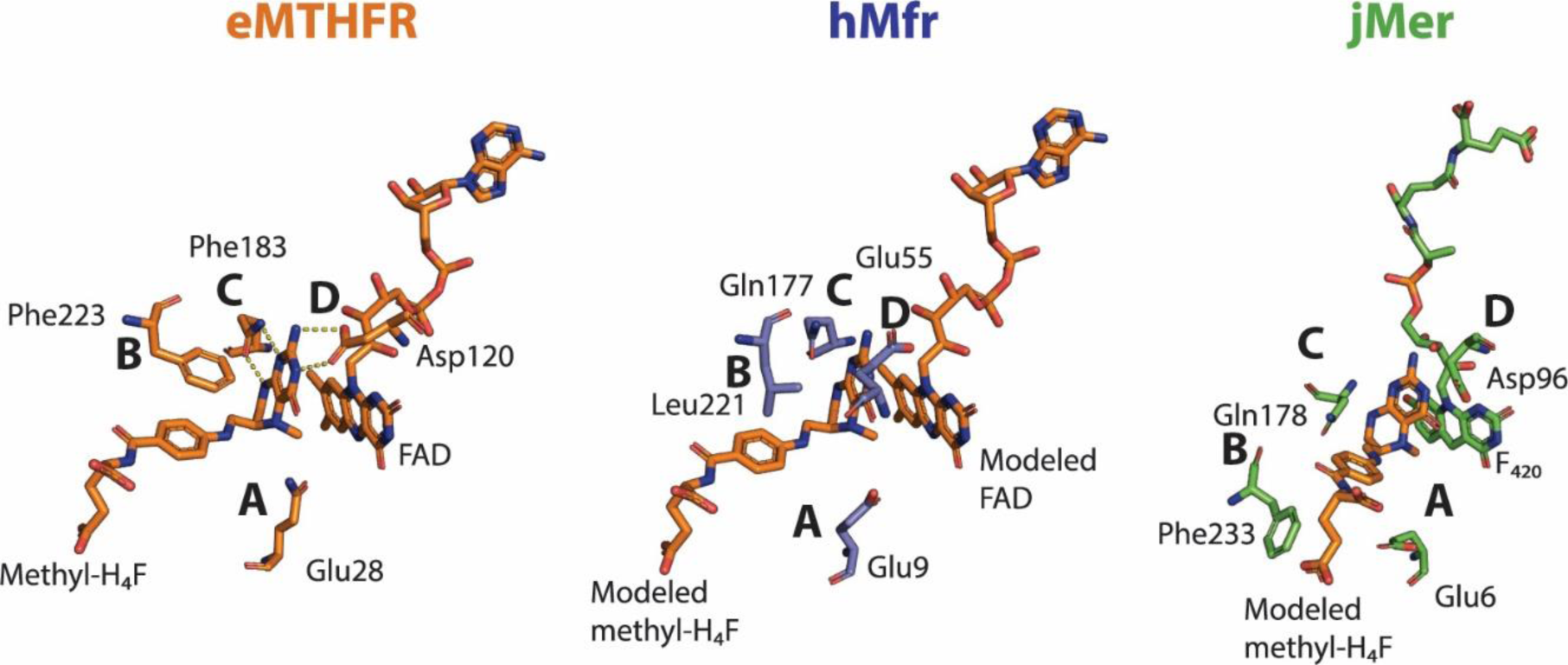
Comparison of the tetrahydropterin binding sites of the crystal structure of eMTHFR and the ternary complex models of hMfr and jMer. Residues from the structure of eMTHFR are colored orange at the carbon atoms, residues from the structure of hMfr are colored blue, and residues from the structure of jMer in complex with F_420_ are colored green. Equivalent positions are indicated by letters.

### Mutational analysis of the C_1_ carrier binding site

The similar amino-acid residues in the active sites of the three methylene-tetrahydropterin reductases indicate that the active site geometries could be similar, despite the fact that these three enzymes share only a very limited degree of amino acid sequence identity. To test whether the amino-acid residues at position A, B, C and D are involved in binding of the C_1_ carrier and catalysis, systematic mutational analyses were performed and the kinetic data were compared with those for the eMTHFR mutants reported in the literature (Table 2).

Position A contains a glutamate residue in all three reductases and replacement with a glutamine residue results in a large decrease in *k_cat_* without a significant change in the *K_m_* values for the C_1_ carrier. For eMTHFR, the *k_cat_* value of the Glu28Gln mutant is reduced from 132 min^-1^ to 0.012 min^-1^ (0.01% residual activity). It has been proposed that Glu28 is the key catalytic residue involved in the protonation of methylene-H_4_F to activate the C_1_ unit for the reaction [11]. In hMfr, the equivalent Glu9Gln mutant showed a decrease from about 594 min^-1^ to 1.3 min^-1^ (0.2%) (Gehl et al. 2023), and the Glu6Gln mutant of jMer showed a decrease from around 18200 min^-1^ to 71 min^-1^ (0.4%). Neither mutant showed a significant increase in the *K_m_* for their respective C_1_ carrier, suggesting that glutamate and glutamine bind in a related manner. X-ray structure analysis of the jMer_E6Q and hMfr_E9Q (Table S1) confirmed an unchanged active site architectures compared with the wild-type enzymes (Figure S3).

Position B in eMTHFR is occupied by Phe223 whose side chain forms π-π interactions with the phenyl ring of methyl-H_4_F. Mutation of Phe223 to alanine or leucine in eMTHFR is associated with a dramatic increase in the *K_m_* values for methylene-H_4_F and NADH without substantially changing the *k_cat_* values [30]. jMer carries a conserved Phe233 at position B but the exchange to alanine or leucine changed the *K*_m_ values only slightly, while the change of the *k_cat_* value (7−73%) was similar to those observed in the eMTHFR mutants (28−135%). The slight increase of the *K*_m_ values in the Phe233 mutants in jMer might indicate that Phe233 is also involved in the binding of the C_1_ carrier, although a strong contribution of π-π interactions as in the case of eMTHFR cannot be concluded. The approximately 6-fold higher *k_cat_* of jMer_Phe233Leu compared to that of jMer_Phe233Ala suggests that the volume of the side chain at this position is important. In hMfr, position B is occupied by the strictly conserved Leu221. Exchange of Leu221 to phenylalanine or alanine increased the *K_m_* values of methylene-H_4_F 2−4 fold, which is again to a much lower extent than those observed in the eMTHFR mutants. In the hMfr_Leu221 mutants, the *k_cat_* value decreased slightly as observed in the case of the equivalent mutants of eMTHFR and jMer. This result supports the hypothesis that Leu221 is involved in the binding of methylene-H_4_F but the strength of interactions to the phenyl ring of the C1 carriers are different in jMer and hMfr compared with those in eMTHFR.

Position C in the ternary complex structure of eMTHFR is occupied by Gln183, whose side chain carboxamide forms a bidentate hydrogen bond with NH8 and N1 [12]. eMTHFR_Gln183Glu and Gln183Ala mutations are associated with a large increase in *K_m_* values for methylene-H_4_F (> 250 fold). In contrast, the Gln183Glu mutation caused no change of the *k*_cat_ value [31]. The exchange of Gln177 in hMfr to glutamate or alanine increased the *K_m_* values for methylene-H_4_F (2-3 fold), but the magnitude was much lower than the case of eMTHFR and the *k_cat_* values significantly decreased (9-17%). Notably, the *K_m_* value for NADH increased in hMfr_Gln177Glu (8 fold), indicating that the acidic side chain also has some effect on the binding of NADH. In contrast, the replacement of Gln178 in jMer did not increase the *K_m_* values for methylene-H_4_MPT, but caused rather a slight decrease (30-40%), while the *k_cat_* value was strongly reduced (0.02-2%). Although H_4_F and H_4_MPT share NH8 and N1 of the pterin ring the interactions between the residue in position C and the pterin ring significantly differ between jMer and eMTHFR. In the jMer_Gln178 mutants, the *K_m_* values for F_420_ substantially increased (6-13 fold) with concomitant decrease of *k*_cat_ (0.1-7%). Since the backbone of Gln178 is part of the F_420_ binding site in jMer, the Gln178 mutation affects the architecture of the active site including the binding of F_420_ rather than the binding of methylene-H_4_MPT.

At position D, the carboxy group of Asp120 forms a bidentate hydrogen bond to N3 and the exo-NH2 group of methyl-H_4_F in the ternary complex of eMTHFR [12]. The mutation of Asp120 to asparagine resulted in a large increase (> 40 fold) of the *K_m_* values for the C_1_ carrier in eMTHFR and a large decrease to 0.3% of the *k*_cat_ value [32]. In the structure of hMfr, a glutamate rather than aspartate residue is located at the equivalent position. The hMfr_Glu55Gln variant did not show a significant increase in the *K_m_* value for methylene-H_4_F but a decrease in the catalytic constant to 30%. In the case of jMer_Asp96Asn, the *K*_m_ value was increased 2.5-fold and the *k*_cat_ was decreased to 4%. These results indicate that the function of aspartate at position D in eMTHFR is not shared with the other two reductases in binding of the C_1_ carrier. For hMfr, Glu55 is not strictly conserved and alanine and valine are found at this position in addition to glutamate and aspartate, which also supports this conclusion.

Based on the mutational analysis at position A, it was possible to extend the proposed model of the catalytic mechanism of MTHFR [11] not only to Mfr (Gehl et al. 2023) but also to Mer. The mechanism is initiated by the protonation of the C_1_ carrier to activate this molecule, followed by a hydride transfer from the hydride donor to the activated C_1_ unit (Figure 7). As the methylene group in methylene-H_4_F and methylene-H_4_MPT is chemically a rather unreactive aminal, it is unlikely that a hydride can be directly transferred. Therefore, the formation of a positively charged 5-iminium cation is crucial for catalysis. The positive charge can be delocalized via the pterin ring system, which stabilizes this intermediate state. In contrast to the uncharged methylene group, the positively charged 5-iminium cation is a good hydride acceptor, which can be transferred from either FADH_2_ in MTHFR, F_420_H_2_ in Mer or NADH in Mfr. On the other hand, even though some identical or similar residues were observed in the three methylene-tetrahydropterin reductases at the predicted binding site of the C_1_ carriers, the effect of mutations did not show common features for the binding of the C_1_ carriers. This finding suggests that the position of the key glutamate residues for the 5-iminium cation formation to the C_1_ carrier is conserved. However, the coordination of the C_1_ carrier in the three reductases is not identical.

### Mutational analysis of the non-prolyl *cis*-peptide bond

Non-prolyl *cis*-peptide (NPCP) bonds are rare in proteins, but they play an important role when they do occur [33,34]. The NPCP bond in jMer is located in the loop after β3, below the central pyridine ring of F_420_, between Gly61 and Val62 (Figure 6). This NPCP bond is conserved at the equivalent position in all known Mer structures and even in other, but not all, enzymes of the bacterial luciferase superfamily, e.g. in the F_420_-dependent glucose-6-phosphate dehydrogenase [35] and the F_420_-dependent alcohol dehydrogenase [36]. It has been proposed that the NPCP bond acts as a backstop for the placement of F_420_ in the active site [19]. In the present study, the role of this NPCP bond was tested by exchanging Val62 to Pro62, presumably resulting in conversion from the NPCP bond to a prolyl-*cis* peptide (PCP) bond. The specific activity of the purified jMer_Val62Pro mutant enzyme was 0.09 U/mg that is much lower than wild-type jMer (160 U/mg) under standard assay conditions.

**Figure 6:**
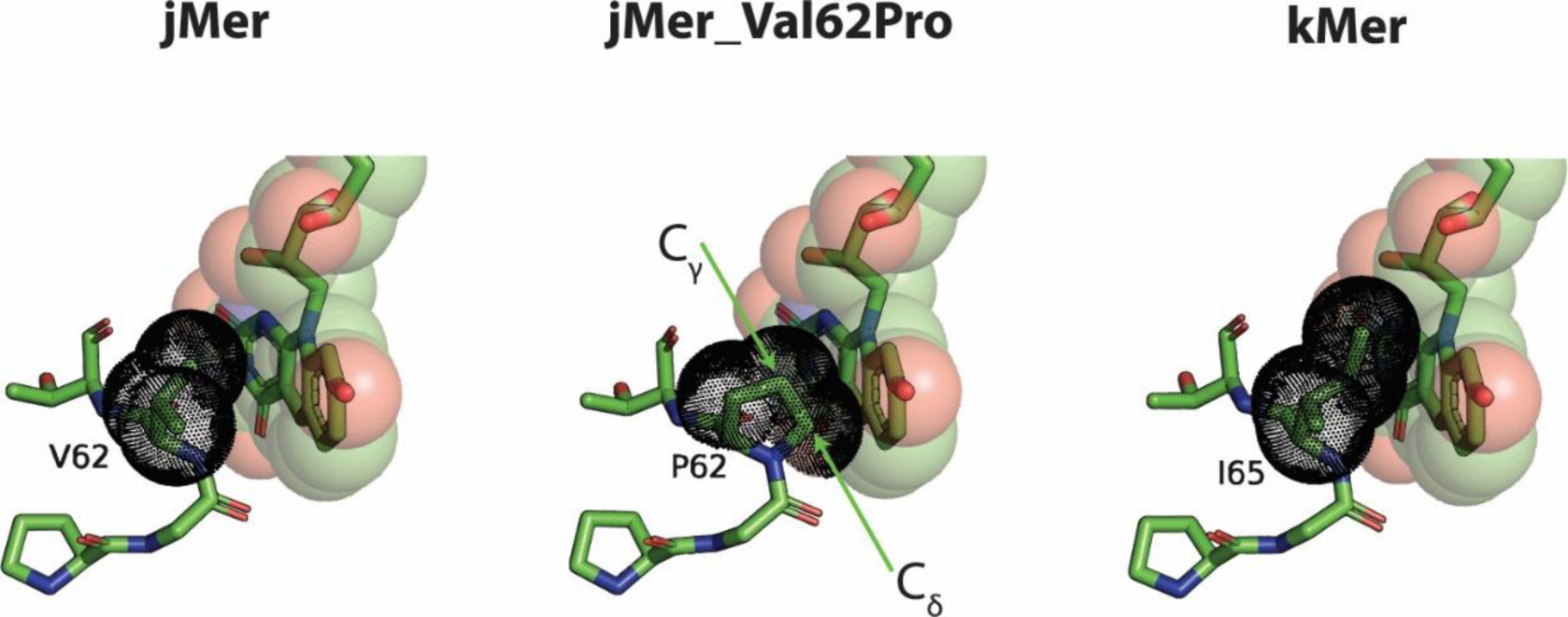
Predicted structural variations of the non-prolyl *cis*-peptide (NPCP) bond region of Mer. F_420_ and the amino acids are shown as stick models with green carbons. The van der Waals radii are shown as transparent spheres for F_420_ and as black dotted spheres for the *cis*-peptide bond regions. The jMer wild type in the binary complex with F_420_ (jMer), the AlphaFold model of jMer_Val62Pro, which contains prolyl *cis*-peptide (PCP) and F_420_, and the crystal structure of Mer from *M. kandleri* (kMer) are shown. F_420_ in kMer was modeled by alignment of the whole protein with the jMer structure in complex with F_420_. The NPCP bond of kMer is placed between Gly64 and Ile65. The side chain of isoleucine in kMer does not overlap with modeled F_420_, whereas the side chain of proline in jMer_Val62Pro is too bulky and collides with F_420_. The position of Cβ, Cγ and Cδ atoms of Val62, Pro62 and Ile65 in jMer, jMer_Val62Pro, and kMer is shown.

**Figure 7:**
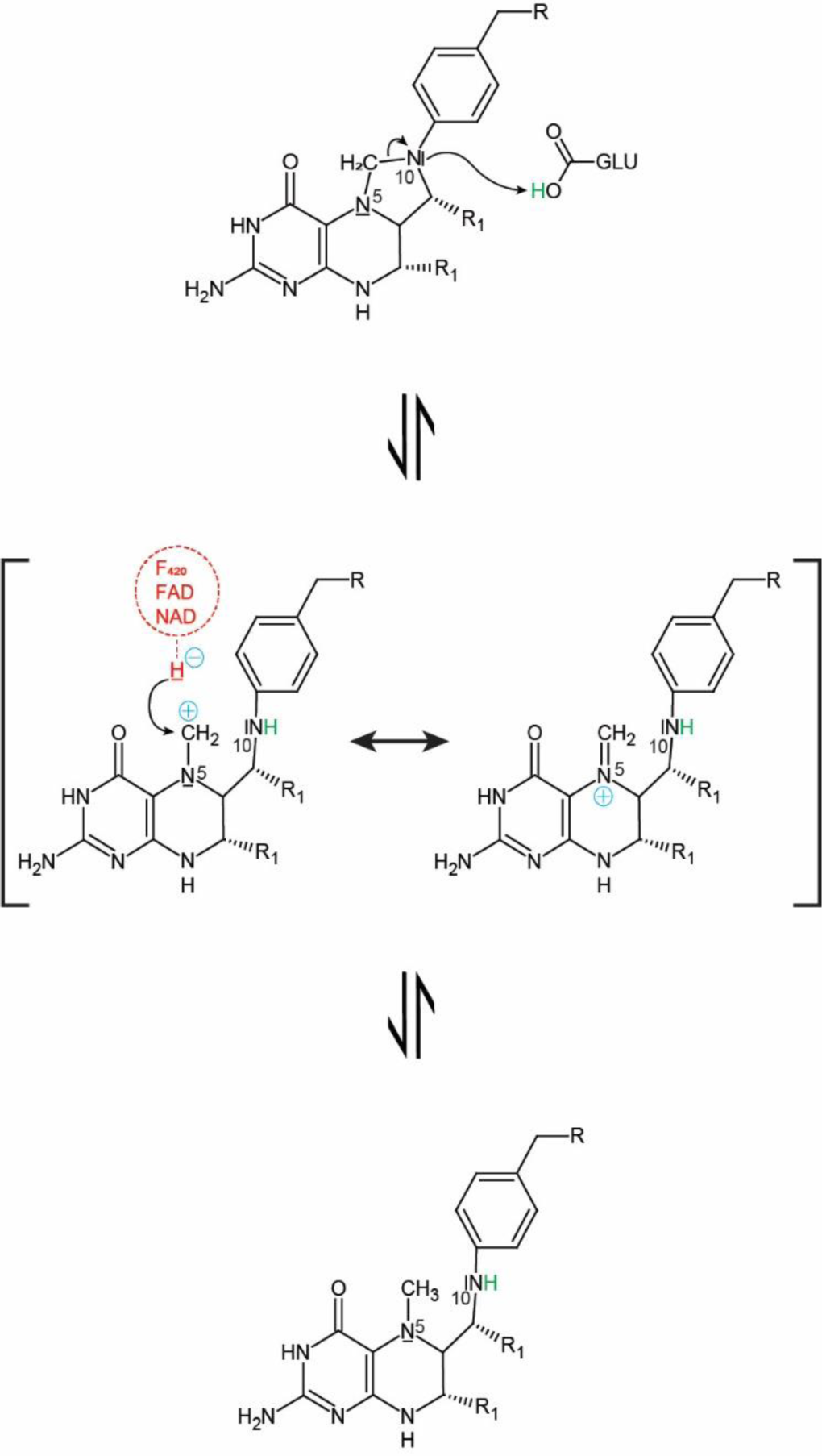
Postulated common catalytic mechanism for all methylene-tetrahydropterin reductases. In the first step, N10 of the pterin part of the C_1_ carrier is protonated leading to the formation of an iminium cation. The positive charge of the iminium cation is delocalized over the pterin ring system, presumably stabilizing the intermediate state. The positively charged iminium cation is an excellent hydride acceptor. After hydride transfer, the reaction product is obtained.

The AlphaFold model of jMer_Val62Pro supported the predicted PCP bond by a very high pLDDT of over 98 in the region (P60 – T63) (Figure 6). Alignment of the jMer wild type binary complex structure and the jMer_Val62Pro model showed that the backbone of the corresponding loop and the C_β_ atoms of Val62 and Pro62 remain in the same position after mutation (Figure 6). However, C_γ_ and C_δ_ of Pro62 occupy the space of F_420_ in the conformation of the wild type enzyme. Although the backstop function of the cis-peptide conformation is maintained, the ability to bind F_420_ is thereby substantially reduced. Spatial consideration indicated that hydrophobic residues of moderate size can replace Val62. Indeed, valine is not strictly conserved at this position and isoleucine is found in other Mer enzymes, e.g. in Mer of *M. kandleri* (Figure 6) [18]. Larger side chains cannot be placed at this position because they would interfere with the loop after α4, which is involved in the binding of the first two hydroxy groups of the F_420_ tail region. Small side chains, such as in alanine, do not reach F_420_ and cannot exert pressure to adjust the conformation of F_420_.

### Phylogenetic analyses of methylene-tetrahydropterin reductases

The presented results indicated that despite minor sequence identity of the three methylene-tetrahydropterin reductases, the (βα)_8_ barrel overall fold, the basal active sites structure, and the key catalytic glutamate residues are conserved. This finding raises the question of whether these three methylene-tetrahydropterin reductases evolved from one common ancestor. From the mechanistic point of view both Mer and Mfr share a one-step hydride transfer process suggesting a closer phylogenetic relationship. On the other hand, structural alignment clearly showed a stronger connection between MTHFR and Mfr. To obtain further insights into the phylogenetic relationship of the three methylene-tetrahydropterin reductases, a phylogenetic tree was constructed using sequences from the bacterial luciferase family including Mer (Mer superfamily; Table S2), the FAD-dependent methylene-H_4_F reductase superfamily (MTHFR superfamily; Table S3), and Mfr for BLAST searches.

The constructed phylogenetic tree shows the clustering of the protein families (Figure 8). Mer is placed in a direct relationship with bacterial luciferases and F_420_-dependent dehydrogenases as previously reported [37]. The MTHFR cluster, consisting of prokaryotic and eukaryotic FAD-dependent MTHFRs, and the mycobacterial Mfr cluster are isolated in the tree with no reliable connection to any other clusters. The tree is therefore an indication that Mfr, Mer and MTHFR are not directly evolutionally related. However, the isolation of the MTHFR and Mfr clusters in the tree may result from missing links due to the bias of the used dataset towards the bacterial luciferase superfamily, as the sequences of Mfr are very limited and no Mfr candidates have been investigated outside the order *Mycobacteriales*. Based on the phylogenic tree, the bottom of the bacterial luciferase family is not occupied with a Mer-like methylene-H_4_MPT reductase. This is an argument favoring the evolution of Mer from an ancestor, whose function was not the reduction of methylene-H_4_MPT.

**Figure 8:**
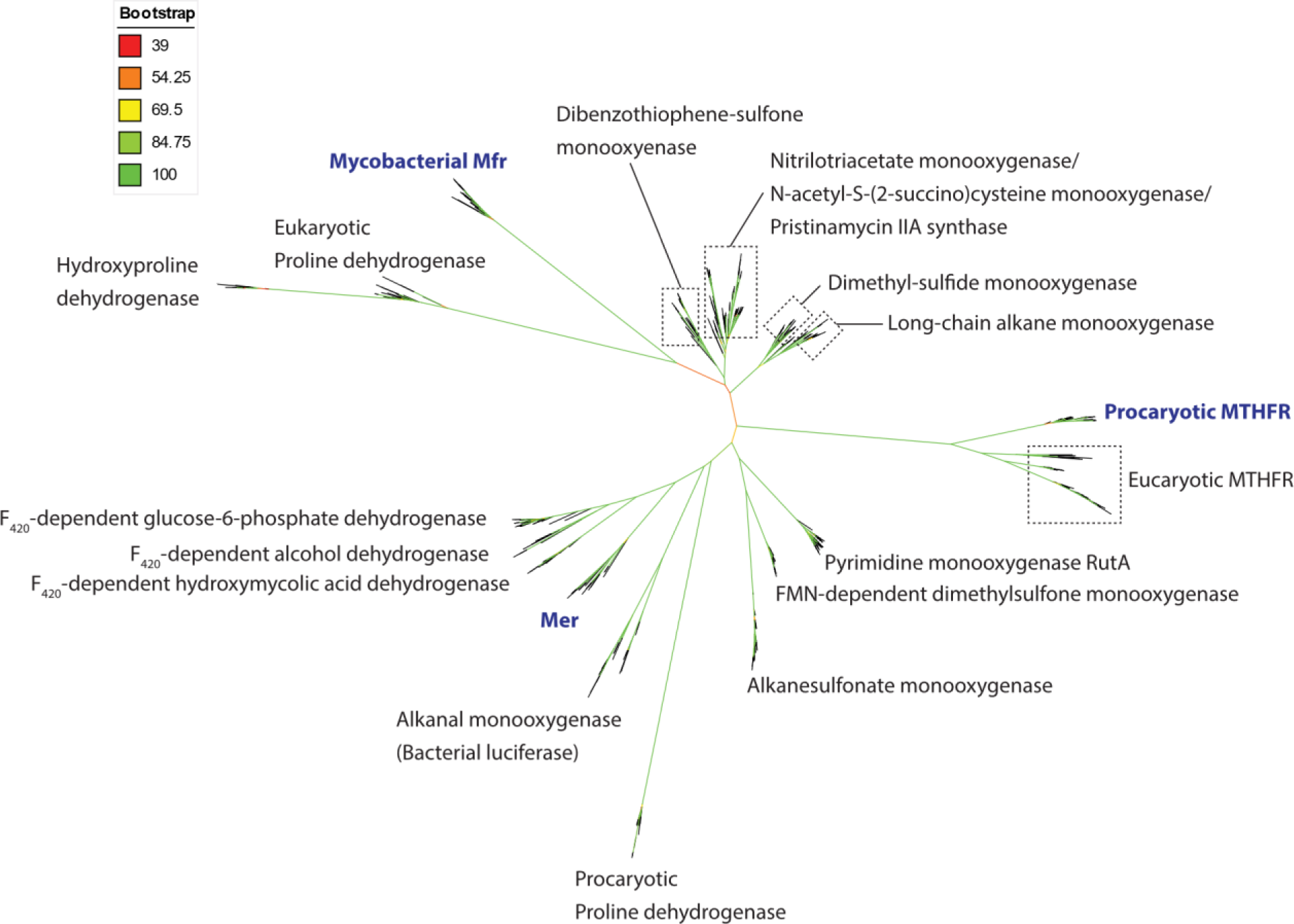
Unrooted phylogenetic tree of bacterial luciferase superfamily members, flavin-linked oxidoreductase superfamily members and Mfr members. The bootstrap values are indicated by a color scheme. The central part of the tree does not allow conclusions about the basal relationship between the tree superfamilies.

The evolutionary relationship between the C_1_ carriers, H_4_MPT and H_4_F, and the whole pathway of C_1_-unit reduction may help to resolve this issue. Biosynthesis of H_4_MPT and H_4_F has been considered to be unrelated [1,38,39] and the enzymes catalyzing the reduction series for C_1_ units bound to H_4_F and H_4_MPT are also considered to be non-homologous [39,40]. This favors Mer as non-homologous to MTHFRs and Mfr. Moreover, eukaryotic and prokaryotic MTHFRs originating from pathways widespread in all domains of life reliably cluster together, which supports the hypothesis of the flavin-dependency of the MTHFR-like enzyme in the last universal common ancestor. The pressing question is why most organisms carry an FAD-dependent MTHFR, while there seems to be no obvious benefit from this prosthetic group because simpler Mfr without flavin can catalyze the same reaction. The function of the ancestral flavin-dependent MTHFR may have been different from what is known. The hypothetical ancestral enzyme reaction may be conserved in any of the homologous enzymes that have not yet been characterized. Researches on the physiological reaction of acetogenic FMN-dependent MTHFRs may resolve this problem. From this hypothetical ancestral flavin dependent enzyme, FAD-dependent MTHFR may have evolved, retaining FAD as an evolutionary relict. If Mfr was evolved from MTHFR, the flavin was then replaced by NADH in mycobacteria later on. To further understand the phylogenetic relationship of the three methylene-tetrahydropterin reductase, biochemical investigations on Mfr functional homologs is important.

The performed structural, mutational and phylogenetic studies on eMTHFR, hMfr and jMer revealed striking similarities in the binding sites of methylene-tetrahydropterin although the specific functions of tested residues essential for the binding of the active part of the C_1_ carrier are not identical. This finding supports an independent origin, from which the same enzymatic activity with a related catalytic mechanism was developed. However, the found similar effects of equivalent mutations in hMfr and eMTHFR rather than in jMer and hMfr or in jMer and eMTHFR keep the hypothesis alive that Mfr has evolved from flavin-dependent MTHFR.

## MATERIAL AND METHODS

### Purification of H_4_MPT and F_420_

Approximately 130 g of *M. marburgensis* cells cultured under nickel-sufficient conditions were used to purify H_4_MPT as previously described [41]. For the purification of F_420_, three cell pellets resulting from the H_4_MPT purification were combined and diluted 1:4 in anaerobic water. The suspension was sonicated using a SONOPLUS GM200 (Bandelin) with a VS-70-T tip attached at 80% power of 100 W for the whole period of 15 min. The suspension was mixed 1:1 with 100% acetone (−20 °C) and stirred in an ice bath for 30 min. The mixture was centrifuged on at 13 000 × g and 4 °C for 20 min. The supernatant was collected and the extraction was repeated twice by adding 50% (v/v) aqueous acetone cooled to −20 °C in a 1:1 ratio regarding the weight of cell debris. The combined supernatants were evaporated at 4 °C until the volume has decreased by at least half. The evaporated solution was centrifuged as described above and applied to a QAE Sephadex A25 column (GE Healthcare) equilibrated with 500 ml of 50 mM Tris/HCl pH 7.5. The column was washed with 500 ml 50 mM Tris/HCl pH 7.5 and then with 500 ml 300 mM formic acid in water. Elution was performed with a single step gradient of 500 ml 50 mM HCl in water. The F_420_ containing fractions were combined and concentrated by evaporation. The concentrate was desalted using a Sephadex G-10 column (Cytiva Life Sciences) equilibrated with water. The F_420_ solution was stored in aliquots at −20°C.

### Mutagenesis and heterologous overproduction of jMer

The vector pT7-7_jMer containing the jMer encoding gene *MJ1534* was used for targeted mutagenesis. The degenerated primers for mutagenesis were designed using NEBaseChanger. A PCR was performed using 10 ng pT7-7_jMer as template, 0.5 µM degenerate primers (Table S4), 1× Q5 reaction buffer (New England Biolabs), 200 µM dNTPs (Thermo Fisher Scientific) and 0.02 U/µl Q5 High-Fidelity DNA Polymerase (New England Biolabs) in a 50 µl reaction volume. Thermocycling conditions were selected according to the manufacturer’s recommendations and the annealing temperature was used as recommended by the NEBaseChanger (Table S4). After PCR, the template DNA was digested with DPNI at 37 °C for 1 hour. The preparation was purified using the NucleoSpin Gel and PCR Clean-up Kit (Macherey-Nagel). The DNA preparation was used for transformation into chemically competent *E. coli* Top10 cells and the cell suspension was plated on agar plates containing 100 µg/ml carbenicillin. After colony formation, 5 ml of LB medium supplemented with 100 µg/ml carbenicillin was inoculated with one colony and grown overnight at 37 °C. The plasmids were isolated using the NucleoSpin Plasmid Kit (Macherey-Nagel) and sequenced by Eurofins using pT7-7_Seq_F and pT7-7_Seq_R as sequencing primers (Table S4). The correct plasmids were transformed into ArcticExpress (DE3) cells and plated on agar plates containing 100 µg/ml carbenicillin and 20 µg/ml gentamicin. One colony was inoculated into 5 ml of LB medium and incubated overnight at 37 °C. A cryo-culture was prepared by mixing 1 ml of 50% glycerol with 1 ml of the overnight culture and flash frozen. Cryo-cultures were stored at −75 °C.

A cryo-culture of *E. coli* ArcticExpress(DE3) containing the desired jMer variant was used to inoculate 100 ml of LB medium supplemented with 100 µg/ml carbenicillin and 20 µg/ml gentamicin. The pre-culture was incubated overnight at 37 °C with shaking at 120 rpm. The main culture contained 2 liters of pre-warmed TB medium supplemented with 100 µg/ml carbenicillin and 20 µg/ml gentamicin and was inoculated with 100 ml of the pre-culture. The main culture was incubated at 37 °C with stirring at 600 rpm until an optical density of 0.6-0.8 was reached. The *mer* gene expression was induced with 1 mM IPTG and the culture was transferred to 21 °C. After 21 h of expression, the culture was harvested by centrifugation at 13 000 × g for 5 minutes at 4 °C. Cells were snap frozen and stored at −20 °C.

### Purification of jMer

For crystallization, approximately 40 g of the *E. coli* cells were suspended in 160 ml of 50 mM Tris/HCl pH 7.5 with 2 mM DTT. The cell suspension was sonicated using a SONOPULS GM200 (Bandelin) with a 50% cycle and 160 W for 5 min per cycle and 5 min pause using a TZ76 tip. A total of 2 cycles were performed. The disrupted cells were fractionated by centrifugation at 30 000 × g for 30 min at 4 °C. The supernatant was heated at 80 °C for 20 min and precipitated proteins were separated by centrifugation at 13 000 × g for 20 min at 4 °C. Ammonium sulfate was added to the supernatant to 60% saturation and the solution was stirred at 4 °C for 20 min. Precipitated proteins were removed by centrifugation at 13 000 × g for 20 min at 4 °C. The supernatant was applied to a Phenyl-Sepharose HP column (15 ml column volume) equilibrated with 50 mM Tris/HCl pH 7.5 containing 2 mM DTT and 2 M ammonium sulfate (buffer A). Buffer B contained 50 mM Tris/HCl pH 7.5 with 2 mM DTT and 10% (v/v) glycerol. The column was washed with 20% buffer B. Elution was performed with a linear gradient from 20% to 100% buffer B in eight column volumes. jMer was eluted from 137 mS/cm to 5 mS/cm conductivity. The corresponding fractions were collected and desalted on a HiPrep G-25 column equilibrated with 50 mM Tris/HCl pH 7.5 containing 2 mM DTT. The desalted solution was applied to a Resource Q column (6 ml column volume) equilibrated with 50 mM Tris/HCl pH 7.5 with 2 mM DTT. jMer was eluted by a linear gradient of 0-250 mM NaCl over fifteen column volumes. The jMer-containing fractions were collected and applied to a HiPrep Sephacryl S-200 HR equilibrated with 50 mM Tris/HCl pH 7.5 containing 150 mM NaCl and 2 mM DTT. The jMer-containing fractions were either used directly for crystallization or, after the addition of 5% (v/v) glycerol, were snap frozen in liquid nitrogen and stored at −75 °C.

A shorter protocol was developed for the characterization of jMer variants. Approximately 10 g of cells were suspended in 40 ml of 50 mM Tris/HCl pH 7.5 containing 2 mM DTT. The cell suspension was sonicated with a TZ73 tip attached to a SONOPULS GM200 (Bandelin) with a 50% cycle and 160 W for 5 min per cycle and 5 min pause. A total of 2 cycles were performed. The disrupted cells were fractionated by centrifugation at 30 000 × g for 30 min at 4 °C. The supernatant was heated at 80 °C for 20 min and precipitated proteins were separated by centrifugation at 13 000 × g for 20 min at 4 °C. The supernatant was diluted 1:1 in 50 mM Tris/HCl pH 7.5 with 2 mM DTT and applied directly to a ResourceQ column (6 ml column volume) equilibrated with 50 mM Tris/HCl pH 7.5 with 2 mM DTT. jMer was eluted by a linear gradient of 0-250 mM NaCl over fifteen column volumes. The jMer-containing fractions were collected and used for the kinetic characterization.

### Activity assay of jMer

Master mixes for the activity assays were prepared in an anaerobic chamber. Brown serum bottles were filled with 100 mM Tris/HCl pH 8.0 supplemented with 10 µM 2-mercaptoethanol, the desired amount of purified F_420_ and H_4_MPT, and 3 mM sodium dithionite. The master mixes were incubated for 15 minutes at 55 °C in a water bath, in which F_420_ was reduced to F_420_H_2_. The enzyme assay was performed in an anaerobic quartz cuvette (1 cm light path) with a final volume of 200 µl. After preheating at 55 °C for 5 min, 15 mM formaldehyde (final concentration) was added, by which methylene-H_4_MPT was generated from H_4_MPT and residual dithionite was quenched. The enzyme reaction was started by the addition of 10 µl of enzyme solution. The reaction was monitored by measuring the increase in absorbance at 401 nm. The catalytic activity was calculated from the extinction coefficient of F_420_ (ε_401_ = 25.9 mM^-1^-cm^-1^). One unit of jMer activity yields one µmole of F_420_ per minute.

### Crystallization and structure determination of jMer

All crystallization experiments were carried out in an anaerobic chamber with a 95%/5% (N_2_/H_2_) atmosphere using the sitting drop vapor diffusion method and 96-well two-drop MRC crystallization plates (Molecular Dimensions). The plates were incubated for one week in the chamber before use. The final protein concentration in each drop was 20 mg/ml. For the drops containing F_420_, a final F_420_ concentration of 2 mM was used. The crystal of the apoenzyme grew in a drop consisting of 35% (v/v) 2-methyl-2,4-pentanediol and 100 mM sodium acetate pH 4.5 and could be frozen directly in liquid nitrogen. The best crystal of the binary complex grew in a drop consisting of 25% (v/v) polyethylene glycol monomethyl ether 550, 100 mM MES pH 6.5 and 10 mM zinc sulfate. Prior to freezing, the crystal was treated with a cryoprotectant solution consisting of the reservoir solution mixed with 20% polyethylene glycol monomethyl ether 550 and F_420_ to a final concentration of 2 mM. A large number of experiments were also carried out using F_420_ in combination with either methylene-H_4_MPT or methyl-H_4_MPT at concentrations ranging from 2 mM to 10 mM substrate concentration in the droplets.

The diffraction experiments were performed at 100 K on the SLS beamline X10SA (Villigen, Switzerland) equipped with a Dectris Eiger2 16 M detector. The data set was processed with XDS and scaled with XSCALE [42]. The phase problem was solved by the molecular replacement method using PHASER [43] with the structure of Mer from *Methanopyrus kandleri* as a search model [18]. The model was built and improved in COOT [44] and refined using Phenix.refine [45] and Refmac [46]. The final model was validated using the MolProbity [47] implementation of Phenix [45]. Data collection, refinement statistics and PDB code for the deposited structure are listed in Table 4.

**Table 3:**
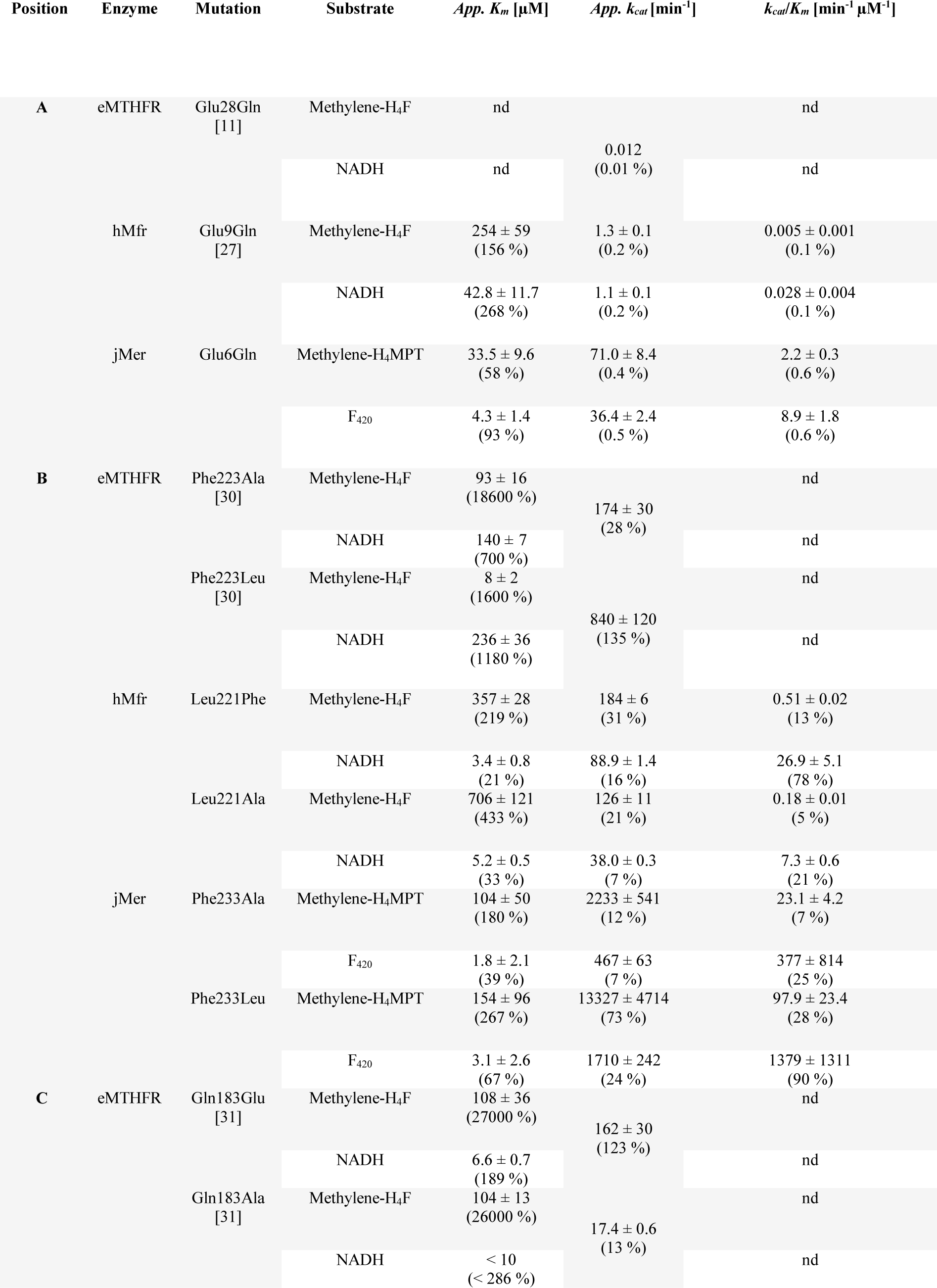
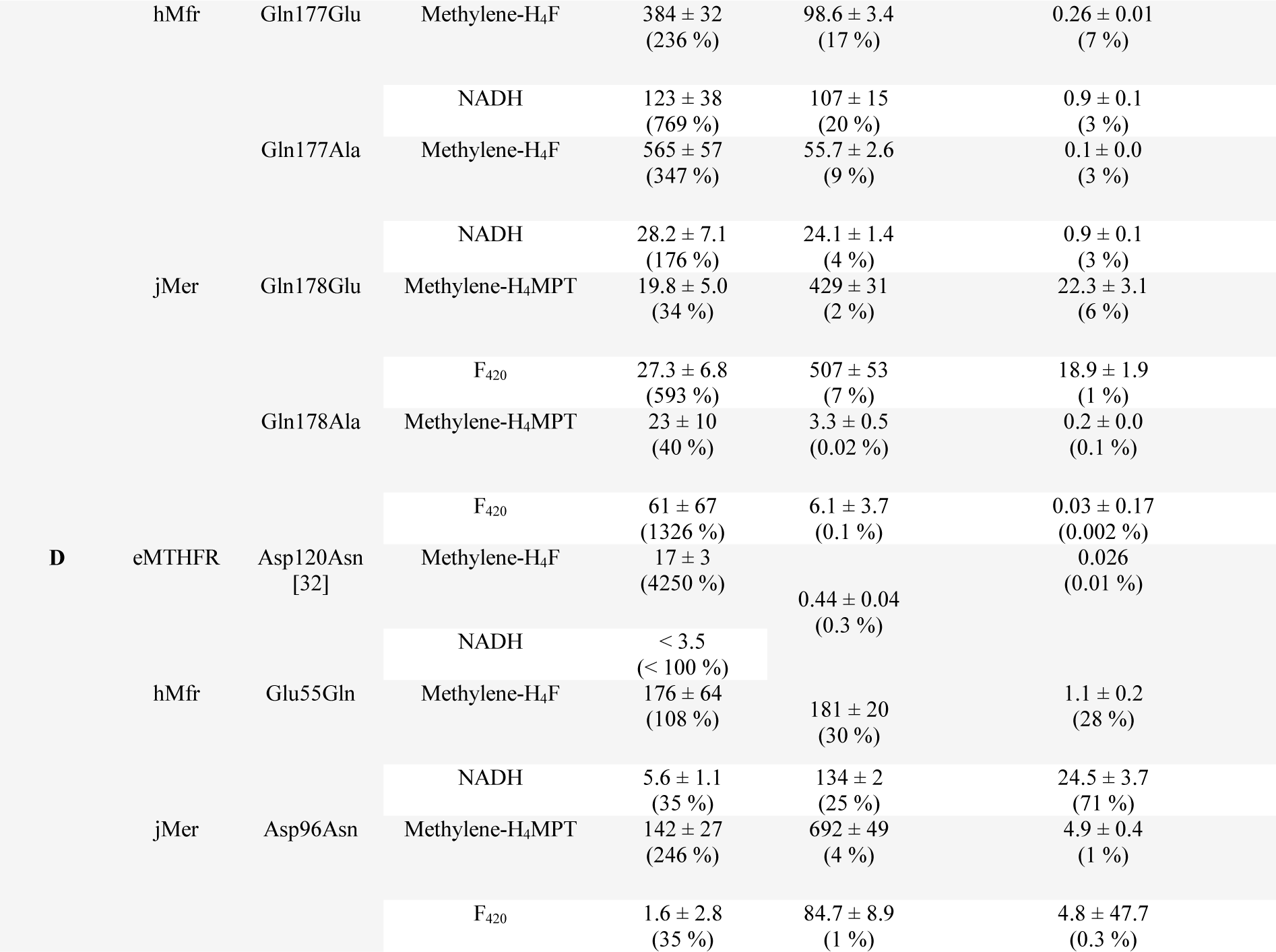
Kinetic constants for the mutants of eMTHFR, hMfr and jMer. References for the eMTHFR mutants and the published hMfr mutants are listed below the mutant name in parenthesis. The relative change in *K_m_*, *k_cat_*, and *k_cat_*/*k_m_* values compared to wild type are in parentheses below the absolute values. nd = not determined.

**Table 4:** Structure determination statistics for jMer apoenzyme and the binary complex of jMer and F_420_.

|  | jMer apoenzyme | jMer + F<sub>420</sub> |
| --- | --- | --- |
| **Resolution range (Å)** | 46.25 - 1.8 (1.864 - 1.8) | 31.39 - 1.902 (1.97 - 1.902) |
| **Space group** | *P* 2<sub>1</sub> 2<sub>1</sub> 2<sub>1</sub> | *P* 4<sub>1</sub> 2<sub>1</sub> 2 |
| **Unit cell dimensions** |  |  |
| a, b, c (Å) | 96.66, 96.28, 166.78 | 95.91, 95.91, 166.02 |
| α, β, γ (°) | 90, 90, 90 | 90, 90, 90 |
| **Unique reflections**<sup>a</sup> | 144123 (14203) | 60825 (5895) |
| **Completeness (%)**<sup>a</sup> | 99.84 (99.54) | 98.83 (97.16) |
| **Wilson B-factor** | 33.65 | 47.64 |
| **Reflections used in refinement** | 144109 (14202) | 60822 (5895) |
| **Reflections used for R-free** | 2002 (200) | 1998 (194) |
| **R-work (%)**<sup>b</sup> | 17.86 (27.11) | 21.95 (32.62) |
| **R-free (%)**<sup>b</sup> | 19.64 (29.09) | 25.25 (33.64) |
| **Protein residues** | 1324 | 657 |
| **RMSD bond lengths (Å)**<sup>c</sup> | 0.008 | 0.014 |
| **RMSD bond angles (°)**<sup>c</sup> | 1.07 | 1.36 |
| **Ramachandran favored (%)** | 97.02 | 96.01 |
| **Ramachandran allowed (%)** | 2.45 | 3.84 |
| **Ramachandran outliers (%)** | 0.54 | 0.15 |
| **Rotamer outliers (%)** | 0.19 | 1.16 |
| **Clash score** | 4.11 | 6.58 |
| **Average B-factor** | 53.52 | 60.38 |

<sup>a</sup> Values relative to the highest resolution shell are within parentheses. <sup>b</sup> R<sub>free</sub> was calculated as the R<sub>work</sub> for 5% of the reflections that were not included in the refinement. <sup>c</sup> rmsd, root mean square deviation.

### Mutagenesis, expression, purification and activity assay of hMfr

hMfr mutants were generated by GenScript. The expression, purification and activity assay procedure was performed as previously described [27].

### Kinetic data processing and figure generation

The graphs and analyses of the kinetic constants were carried out using Python 3.7 with Jupyter Notebook (version 6.1.4) [48] as the development environment and the following packages: os, pandas [49], seaborn [50], matplotlib [51], NumPy [52], SciPy [53]. The code can be found at GitHub (https://github.com/ManuelGehl/Enzyme-kinetic-fitting). The chemical structures were created using ChemSketch and edited using Adobe Illustrator.

### Phylogenetic tree construction

Seed sequences from the bacterial luciferase family (Table S2) and the FAD-linked reductase superfamily (Table S3) together with the amino acid sequence of hMfr were used for separate BLAST searches against the clustered non-redundant protein sequence database. The results were filtered to exclude any cluster with more than 90% sequence identity, and all clusters containing at least 3 members were selected. The query sequences were reinserted into the dataset and sequences marked as partial were removed. A multiple sequence alignment was performed using MUSCLE [54] and a maximum likelihood tree was constructed using IQTree [55]. The best fitting evolutionary model was found to be WAG+F+I+G4 and the ultra-fast bootstrap method [56] was used to incorporate bootstrap values. The tree was visualized using iTOL[57].

## ACKNOWLEDGEMENT

The authors thank Robert H. White for providing us the plasmid for the production of jMer. M.G. thanks the International Max Planck Research School thesis-committee Members: Tobias Erb and Rolf Thauer.

## DATA AVAILABILITY STATEMENT

Data available in article supplementary material.

## SUPPORTING INFORMATION

Additional supporting information can be found online in the Supporting Information section at the end of this article.

## SUPPLEMENTARY INFORMATION

**Table S1:** Structure determination statistics for jMer_E6Q and Mfr_E9Q.

|  | **jMer_E9Q** | **Mfr_E9Q** |
| --- | --- | --- |
| **Resolution range (Å)** | 36.66 - 2.0 (2.072 - 2.0) | 45.8 - 1.75 (1.813 - 1.75) |
| **Space group** | *I* 4<sub>1</sub> 2 2 | *P* 4<sub>3</sub> |
| **Unit cell dimensions**<br>a, b, c (Å)<br>$\alpha$, $\beta$, $\gamma$ (°) | 98, 98, 200.78<br>90, 90, 90 | 45.8, 45.8, 254.6<br>90, 90, 90 |
| **Unique reflections**<sup>a</sup> | 33180 (3259) | 51862 (5183) |
| **Completeness (%)**<sup>a</sup> | 99.07 (98.72) | 98.91 (99.58) |
| **Wilson B-factor** | 46.17 | 25.15 |
| **Reflections used in refinement** | 33124 (3237) | 51850 (5183) |
| **Reflections used for R-free** | 1998 (196) | 1998 (196) |
| **R-work (%)**<sup>b</sup> | 19.29 (39.87) | 19.97 (33.18) |
| **R-free (%)**<sup>b</sup> | 23.44 (43.02) | 23.15 (39.49) |
| **Protein residues** | 331 | 586 |
| **RMSD bond lengths (Å)**<sup>c</sup> | 0.007 | 0.006 |
| **RMSD bond angles (°)**<sup>c</sup> | 0.90 | 0.84 |
| **Ramachandran favored (%)** | 96.96 | 98.79 |
| **Ramachandran allowed (%)** | 2.74 | 1.21 |
| **Ramachandran outliers (%)** | 0.30 | 0.00 |
| **Rotamer outliers (%)** | 0.77 | 0.00 |
| **Clash score** | 6.54 | 3.39 |
| **Average B-factor** | 64.45 | 33.79 |

<sup>a</sup> Values relative to the highest resolution shell are within parentheses. <sup>b</sup> R<sub>free</sub> was calculated as the R<sub>work</sub> for 5% of the reflections that were not included in the refinement. <sup>c</sup> rmsd, root mean square deviation.

**Table S2:**
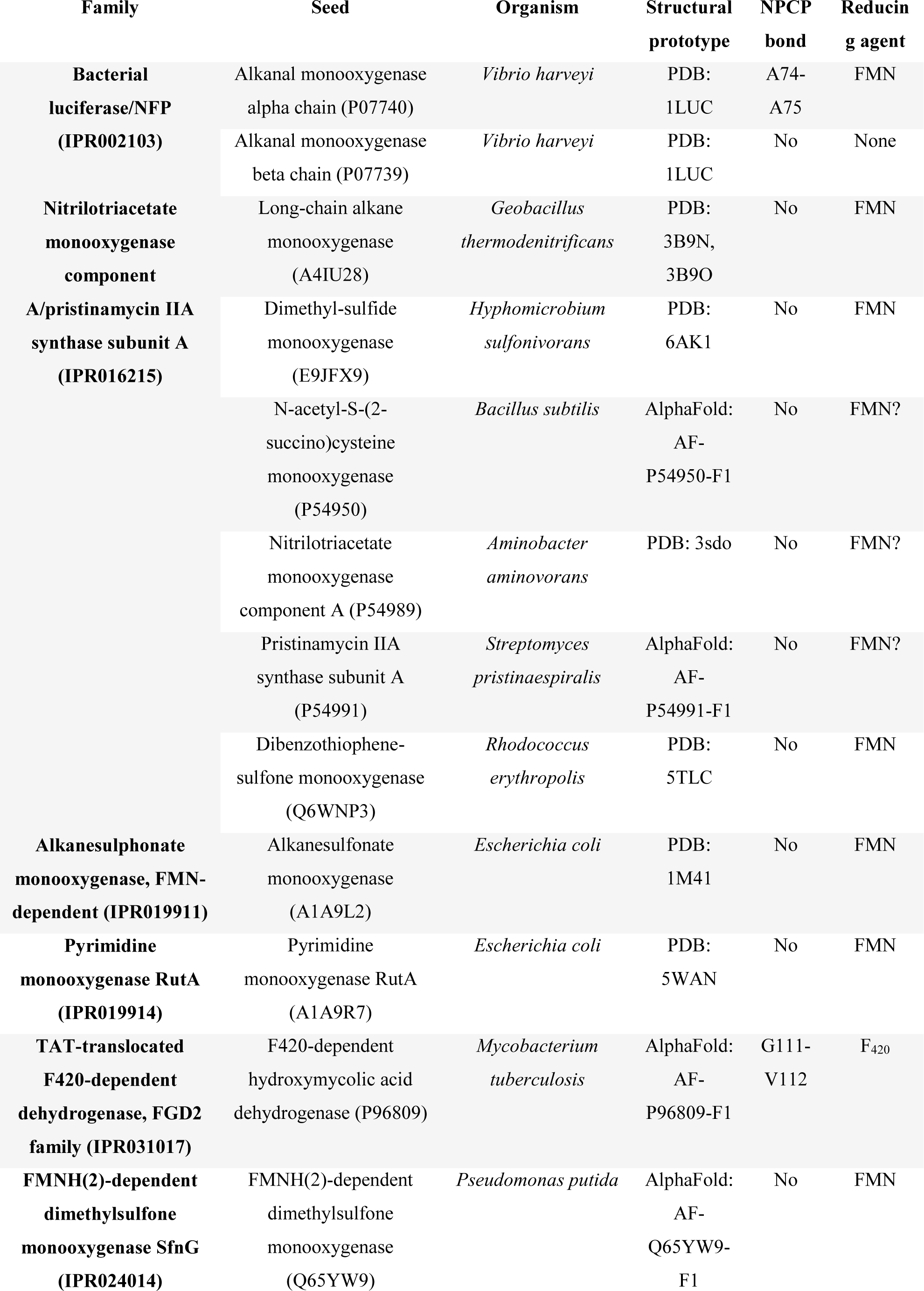
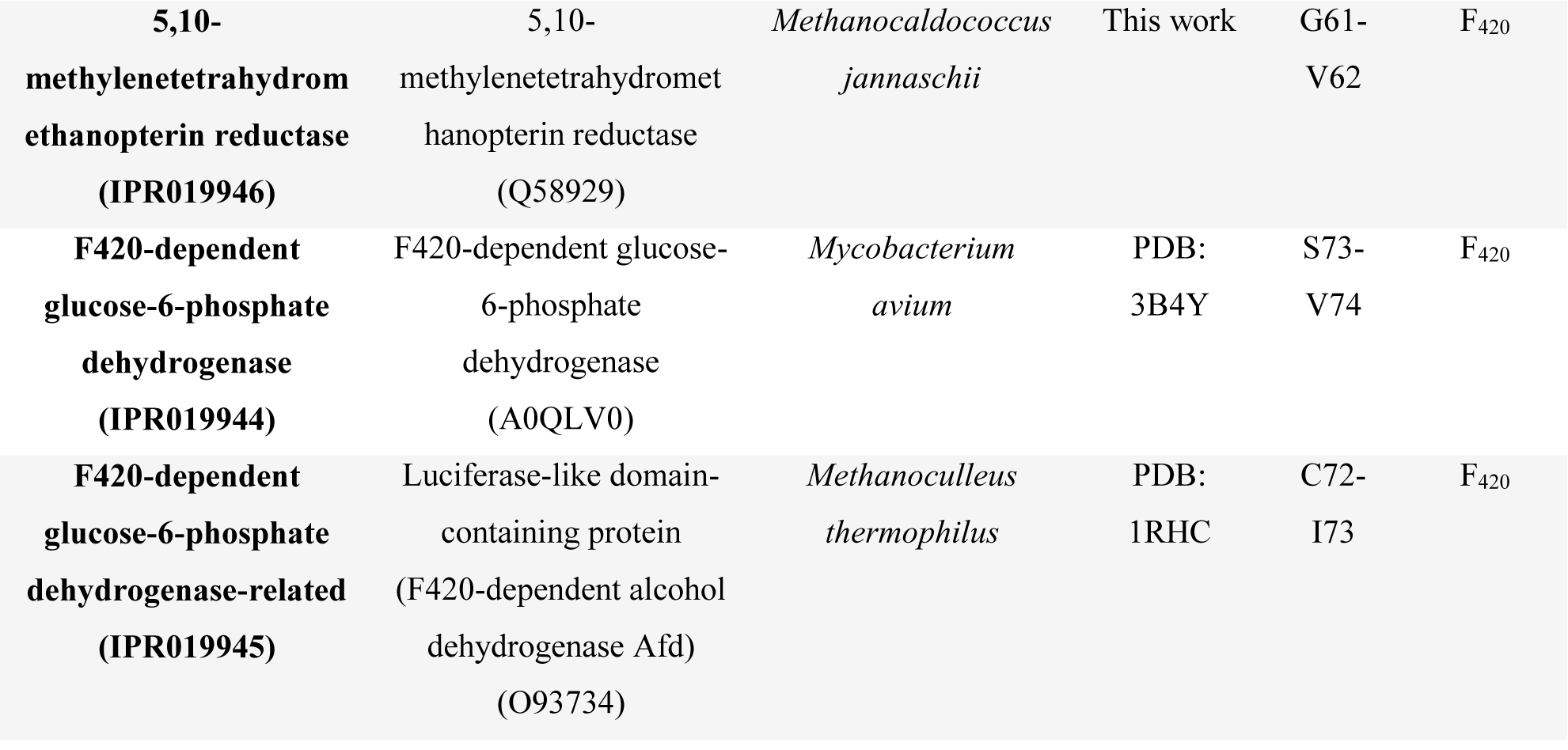
Seed sequences for the Superfamily of Bacterial Luciferases.

**Table S3:**
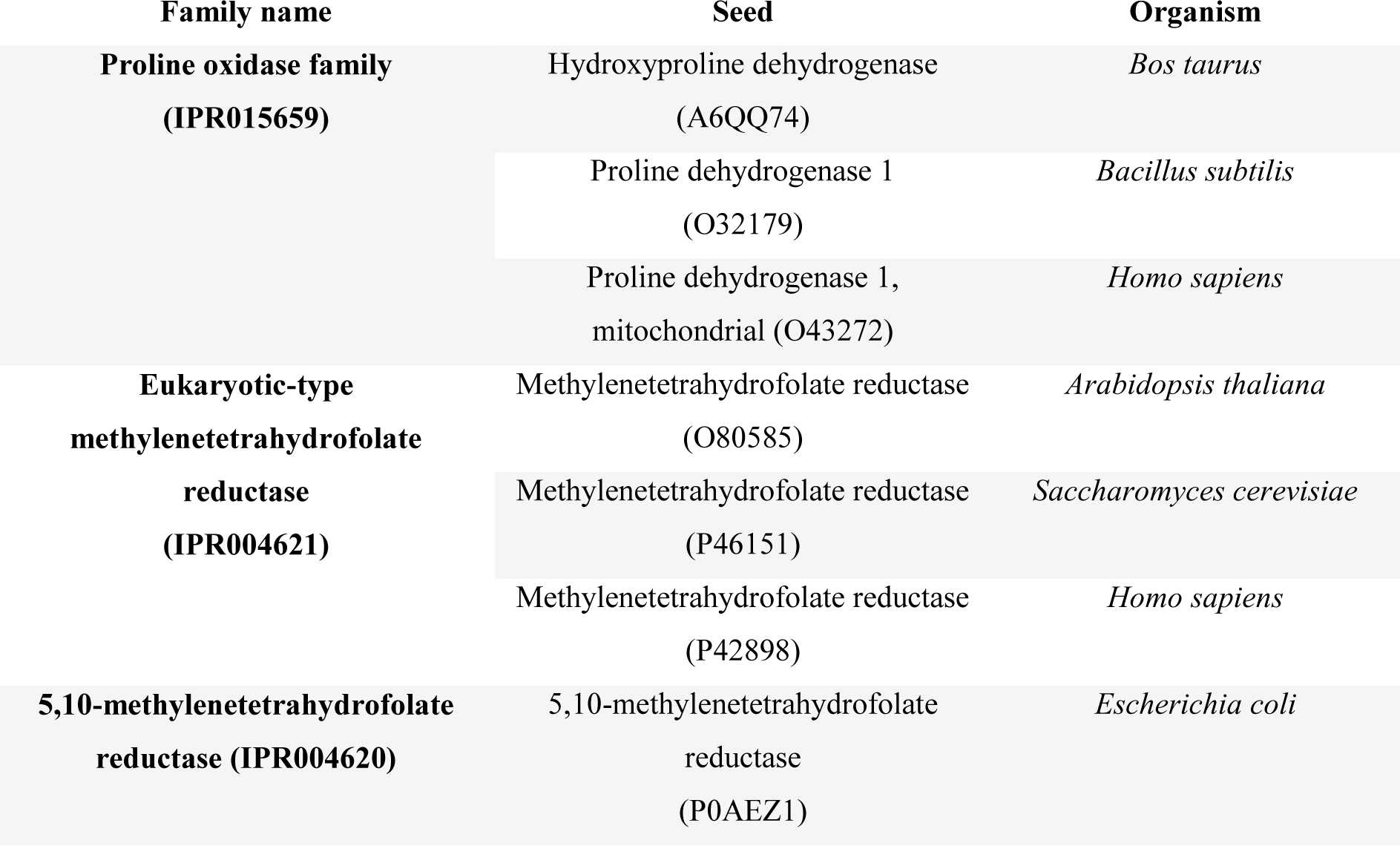
Seed sequences for the Superfamily of FAD-linked reductases.

**Table S4:**
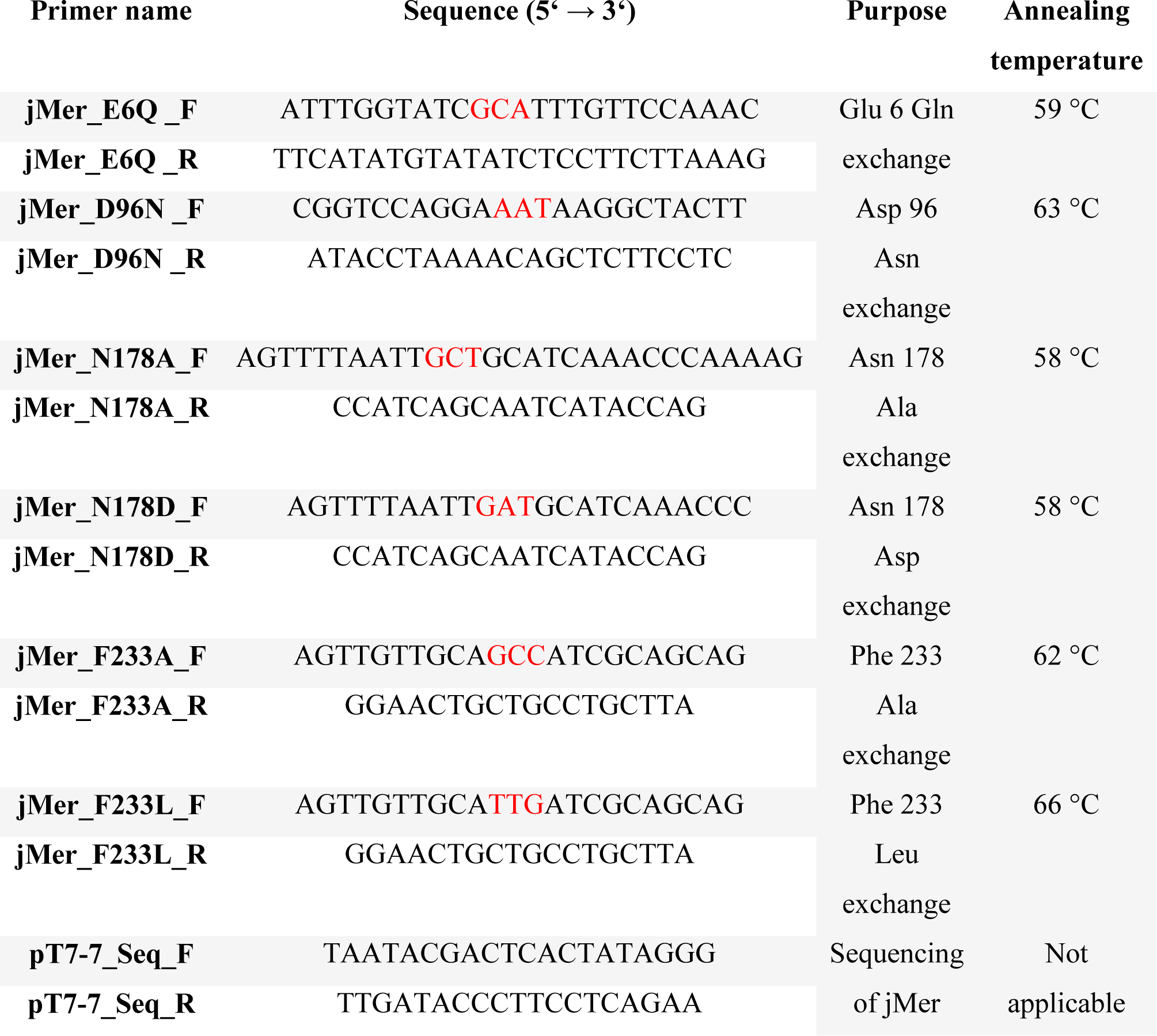
List of primers used for mutagenesis of jMer and sequencing of the ORF of jMer in pT7-7_jMer. The degenerated nucleotides are marked in red.

**Figure S1:**
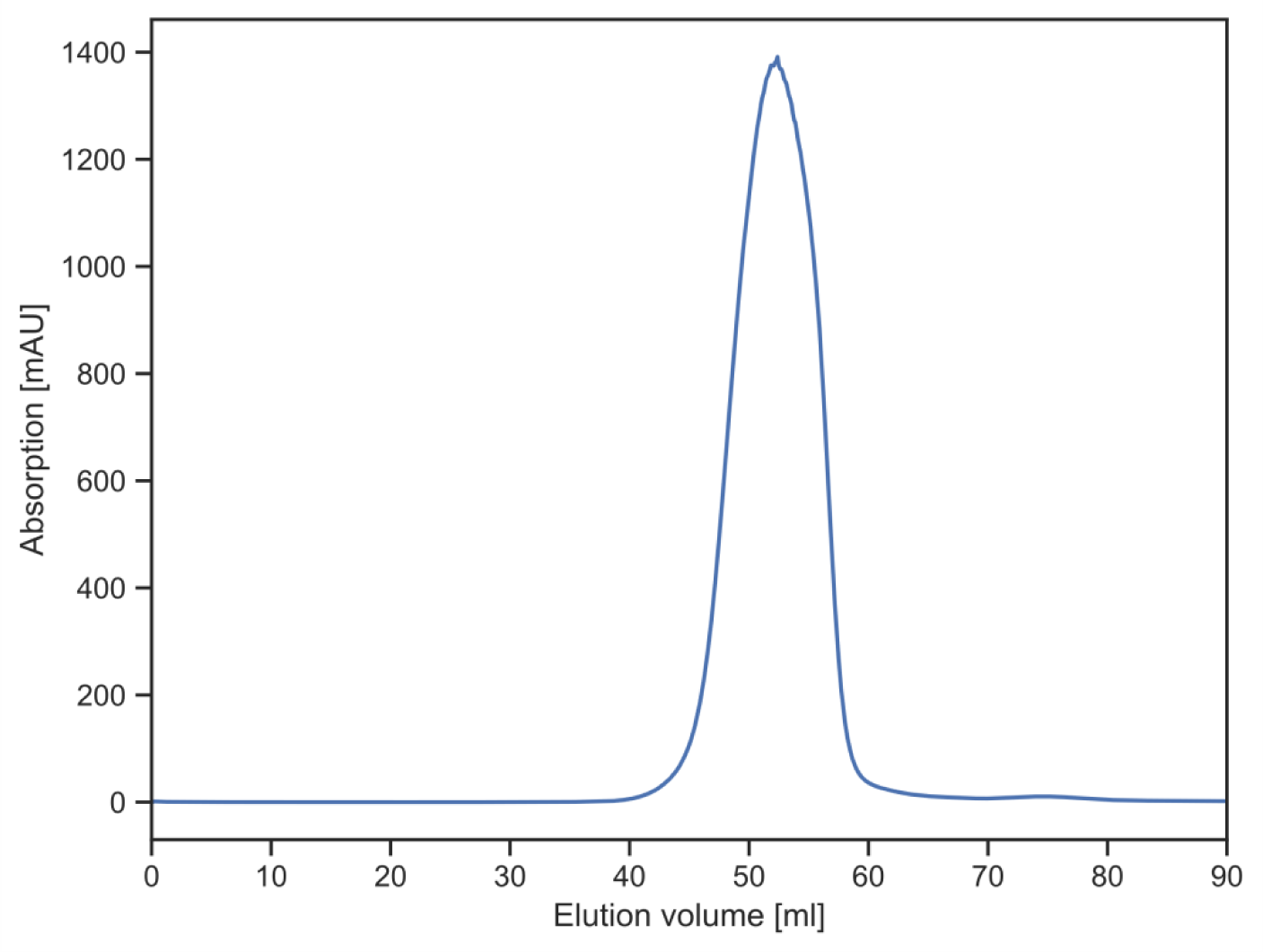
Size-exclusion chromatography of jMer using a HiPrep Sephacryl S-200 HR column. The peak centered around 50 ml elution volume corresponds to jMer and to a molecular mass of approximately 80 kDa.

**Figure S2:**
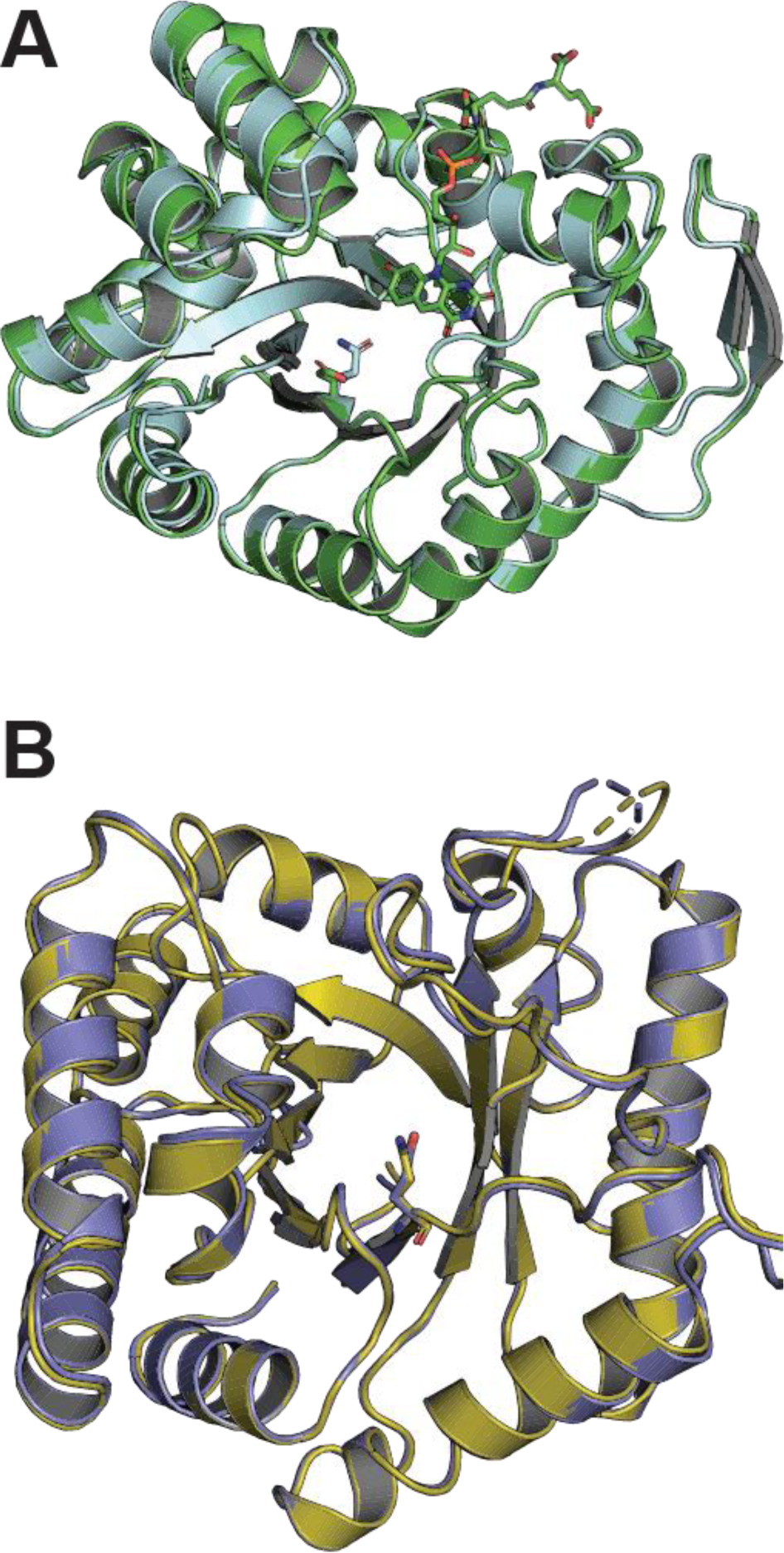
**(A)** Comparison of the structures of jMer wild type (green) and jMer_E6Q (light blue). **(B)** Comparison of the structures of hMfr wild type (dark blue) and hMfr_E9Q (yellow). The glutamate residues are depicted as ball-and-stick model.

